# Understanding the future risk of bat coronavirus spillover into humans – correlating sarbecovirus receptor usage, host range, and antigenicity

**DOI:** 10.1101/2025.09.03.673949

**Authors:** Nazia Thakur, Joseph Newman, Dongchun Ni, Jeffrey Seow, Abigail L Hay, Yeonjae Li, Ayush Upadhyay, Babatunde E Ekundayo, Thomas P Peacock, Kelvin Lau, Katie J Doores, Dalan Bailey

## Abstract

Sarbecoviruses interact with their receptor, angiotensin converting enzyme 2 (ACE2), via the receptor binding domain (RBD) of Spike, the immunodominant target for neutralising antibodies. Understanding the interplay and correlation between ACE2-determined host range and antigenicity is vitally important for understanding the zoonotic potential of related bat sarbecoviruses. Using binding assays, pseudotype-entry assays and a diverse panel of mammalian ACE2 proteins, we examined the host range and related antigenicity of multiple bat coronaviruses. Broad bat ACE2 usage (a *generalist* phenotype) was most common in clade 1 sarbecoviruses, including SARS-CoV-2 and the BANAL isolates from Laos. In contrast, clade 3 (e.g., RhGB07) and 5 (e.g., Rc-o319) sarbecoviruses exhibited more restricted ACE2 usage (a *specialist* phenotype). A novel structure for RhGB07 Spike further helped to identify RBD residues associated with this receptor specialism. Interestingly, the generalist phenotypes were largely maintained with more diverse mammalian receptor libraries, including human, non-human primate, livestock, rodent ACE2 and potential intermediate reservoir hosts (e.g., civet, racoon dog, pangolin), while specialists, like RhGB07, exhibited wider phenotypic diversity. The impact of SARS-CoV-2’s continued evolution in humans was also examined, identifying an expanding and/or shifting pattern of generalism for variants, especially Omicron and its sub-lineages. Furthermore, we compared and correlated these entry phenotypes with antigenicity using sera from SARS-CoV-2 convalescent individuals. Clade 1 viruses, phylogenetically related to SARS-CoV-2, were antigenically the most similar, with robust evidence for cross-neutralisation; however, there was still evidence for limited cross-neutralisation across the entire sub-genus. Finally, using monoclonal antibodies, derived from COVID-19 vaccinees with breakthrough infections, we pin-pointed the antibody epitope classes responsible for wider neutralisation. Our research indicates that generalist ACE2-using sarbecoviruses are phylogenetically and antigenically related to SARS-CoV-2.

## Introduction

Severe respiratory coronavirus 1 (SARS-CoV-1) and more recently SARS-CoV-2, responsible for the COVID-19 pandemic, are both thought to have spilled over from *Rhinolophus* bat species, highlighting the significant risk of bat sarbecoviruses to human and animal health [1, 2]. Our current understanding of the sarbecovirus subgenus is that this group encompasses hundreds of related viruses, exhibiting high genetic diversity arising from recombination and multispecies host spillover events [1–3]. Phylogenetic analyses based on the receptor binding domain (RBD) clusters the extant sequences into five distinct clades: clade Ia which includes SARS-CoV-1, clade Ib which includes SARS-CoV-2, and clades II-V; with the vast majority of sequences coming from bats [4–11]. The angiotensin converting enzyme 2 (ACE2) protein has been identified as the receptor for many of these sarbecoviruses, with the exception of the clade II viruses, for which the receptor remains unknown [12]. To what extent human ACE2 usage is conserved across the sarbecovirus phylogeny is a source of intense investigation [13, 14]. Although RaTG13, isolated from a *Rhinolophus affinis* horseshoe bat, is highly similar to SARS-CoV-2 at the RBD amino acid level, we previously showed it does not share the same broad *generalist* ACE2 usage profile [15]. However, we identified mutations within the RaTG13 RBD-ACE2 interface that could increase human ACE2 (hACE2) usage, which were consistent with the mechanism of action underpinning the early adaptation of SARS-CoV-1 to civet and then human ACE2 [15, 16]. Since 2020, multiple sarbecoviruses have been identified in bat reservoirs that already have high affinity for hACE2, for example, the BANAL viruses found in Laos (BANAL-20-52, BANAL-20-103 etc.) [17], raising intriguing questions about the molecular, ecological and evolutionary drivers for ACE2 generalism in bat reservoirs.

As the first step in the life cycle, receptor engagement is critical in determining virus host-range and species susceptibility. The identification of other human ACE2 using sarbecoviruses with generalist receptor usage profiles [10, 13, 14, 18] is perhaps unsurprising, given the high sequence similarity of some these viruses [17]. However, to date, there have been no highly efficient (equivalent to SARS-CoV-2) human ACE2-using sarbecoviruses identified outside of clade 1. This underpins the need to better characterise the diversity of receptor usage across the whole sub-genera. Furthermore, how this spectrum of generalism and specialism correlates with bat ACE2 usage and bat ecology is also an important area for consideration. A previous receptor usage study using 46 ACE2 orthologues from phylogenetically diverse bat species, showed that 30 of these bat ACE2s were able to support entry of both SARS-CoV-1 and SARS-CoV-2. Further mutational analysis revealed that amino acid residues at positions 27, 31 and 42 in the bat ACE2 were important determinants of bat host-range [19]. RBDs from a more diverse range of sarbecovirus were shown to bind to various *Rhinolophus* ACE2 orthologues with high affinity, in particular clade Ia and Ib viruses [13]. Also, Roelle et al., performing entry assays with chimeric sarbecovirus RBDs in a SARS-CoV-2 Spike backbone, detected entry in cells overexpressing human and *Rhinolophus* ACE2, including for clade III representatives BtKY72 and Khosta-2 [11]. Allied with this is a need to understand how host range and receptor tropism relates to antigenicity. For example, are there human ACE2-using generalist sarbecoviruses that are antigenically totally distinct from SARS-CoV-2? Related, does pre-existing immunity to SARS-CoV-2 protect against, or prevent, related bat sarbecoviruses emerging in the human population? Other studies have shown that monoclonal antibodies derived from macaques immunised with SARS-CoV-2 subunit vaccines are able to neutralise clade I (SARS-CoV-1, WIV-1, SARS-CoV-2, Pangolin-GD) and clade III sarbecoviruses (BtKY72 and PRD-0038) [20]. However, a more detailed analysis of the breadth of COVID-19-derived immunity and the important epitopes that might define cross-protection is lacking. Another important aspect to consider is that over the course of the COVID-19 pandemic, SARS-CoV-2 itself has shifted species tropism [21, 22]. Single amino acid changes in SARS-CoV-2 variant Spikes have expanded its tropism, e.g. to rodents (N501Y) and ferret/mink ACE2 (N501T) [21, 23, 24]. In addition, there are also examples of entirely new animal reservoirs being established, e.g. white-tailed deer, that are able to sustain onward transmission [25–27]. In many cases this can likely be considered coincidental to the continued evolution of SARS-CoV-2 to human ACE2; however, the implications for bat ecology have perhaps been overlooked. Indeed, it is not clear whether novel SARS-CoV-2 variants have evolved enhanced tropism for bat ACE2 proteins. This might be of significance in biodiverse regions such as South America, where anthropogenic spillover (reverse zoonosis) of SARS-CoV-2 into native bat species is a conceivable risk.

To address these questions, we examined the receptor usage of a panel of 15 representative full-length bat sarbecovirus Spike proteins, using ACE2 libraries representing 16 bat species and a further 18 mammalian species of direct anthropogenic and/or zoonotic spillover relevance. We identified that clade Ia and Ib viruses widely, although not exclusively, exhibited generalist bat ACE2 usage, whereas clade III and V viruses were far more specialist, i.e. using only a limited number of receptors. A novel Spike trimer protein structure for one *specialist*, the UK-isolated RhGB07, identified several amino acid motifs that contribute to this host-range tropism. Interestingly, the patterns of generalism and specialism seen with bat receptors were largely mirrored when examining a wider mammalian library, a correlation which highlights that ecological and/or viral drivers of broad receptor usage tropism in bats could potentiate spillover risk into non-bat species. Significantly, bat ACE2 usage tropism has also shifted during the evolution of SARS-CoV-2 in human populations, highlighting underappreciated risks of reverse zoonosis. To broaden our understanding of the wider risk to human populations of future sarbecovirus spillover, we conducted serological assays using COVID-19 convalescent human sera as well as monoclonal antibodies derived from breakthrough infections. Importantly, it appears that all generalist ACE2 users are antigenically related to SARS-CoV-2. Somewhat surprisingly, even phylogenetically distinct, specialist ACE2-using sarbecoviruses still retained some antigenic relatedness to SARS-CoV-2, albeit with lower-level binding and neutralisation titres. This suggests a degree of cross-protection (against other sarbecoviruses emerging in human populations) might be conveyed by pre-existing COVID-19 immunity and outlines the potential of pan-sarbecovirus vaccines.

## Materials and Methods

### Cells

HEK293T cells were used to generate lentiviral pseudoparticles, and BHK-21 cells for receptor screens. HEK293T cells stably expressing human angiotensin converting enzyme 2 (HEK293T-hACE2) were used for pseudotype neutralization assays. Cells were maintained in Dulbecco’s Modified Eagle Medium (DMEM, Sigma-Aldrich) supplemented with 10% FBS (Life Science Production), 1% 100mM sodium pyruvate (ThermoFisher Scientific), and 1% penicillin/streptomycin (10,000Uml−1; Life Technologies) at 37 °C in a humidified atmosphere of 5% CO2. Stable cell lines were generated, as described previously [28], using a lentiviral transduction system under 1 μg/mL puromycin (ThermoFisher Scientific) selection.

### Plasmids

Codon optimised bat sarbecovirus Spike constructs were synthesised and subcloned into pcDNA3.1 (+) with a C-terminal FLAG tag (BioBasic, Canada). Human codon-optimised, Δ19-truncated Spike genes from RhGB07 and RfGB02 were subcloned into pcDNA3.1 (+) [10]. Mammalian ACE2 constructs were subcloned into pDISPLAY with an N-terminal signal peptide (the murine Ig k-chain leader sequence) and HA-tag (BioBasic, Canada; Twist Bioscience, USA). Accession numbers can be found in **S1 and S2 Tables**, respectively. All SARS-CoV-2 variant Spike expression plasmids were based on codon optimised SARS-CoV-2 lineage B (Pango) reference sequence (GenBank ID NC_045512.2), with an additional Δ19 cytoplasmic tail truncation. Mutants of B.1 and BA.1, and RBD chimeras of SARS-CoV-2 and Rs4231 were generated by site-directed mutagenesis, using the QuikChange II Lightning Multi Site-Directed Mutagenesis kit (Agilent).

### Lentiviral pseudotype production

Viral pseudotypes bearing the Spike of different bat sarbecoviruses were generated as described previously [29]. Briefly, HEK293T cells were transfected with p8.91 (gag/pol), CSFLW (lentivirus backbone expressing Firefly luciferase) and the Spike construct of interest using PEI transfection reagent (Merck). Virus was harvested 48 and 72 h post transfection and centrifuged and frozen and −80°C to remove cellular debris. Viruses were titrated on HEK293T-hACE2 stable cell lines, or where viruses used hACE2 poorly, HEK293T cells were transfected with the cognate bat ACE2 receptor (i.e., *Rh. cornutus* for Rc-o319, *Rh. ferrumequinum* for RhGB07 and RfGB02). Luciferase signals were measured 3 days later using Bright-Glo (Promega).

### ACE2 receptor screen

Receptor usage screens were conducted as described previously [15, 21, 29]. BHK-21 cells were seeded in 6-well plates at 7.5 × 10^5^ /well in DMEM-10% 1 day prior to transfection with 500 ng of different species, ACE2-expressing constructs or empty vector (pDISPLAY) (**S2 Table**) in OptiMEM (ThermoFisher Scientific) and TransIT-X2 (Mirus Bio) transfection reagent, according to the manufacturer’s recommendations. The next day, cells were re-plated into white-bottomed 96-well plates at 2×10^4^/well and infected with pseudoparticles equivalent to 10^5^ to 10^6^ relative light units (RLU) for 48 h at 37°C, 5% CO_2_. To quantify firefly luciferase, media was replaced with 50 μL Bright-Glo substrate (Promega) diluted 1:2 with PBS. The plate was incubated in the dark for 2 min and then read on a Glomax Multi+ Detection System (Promega) as above. Comma-separated values (CSV) files were exported for analysis. Biological replicates were performed 3 times.

### Micro virus neutralisation assays

Neutralisation assays were conducted as described previously. COVID-19 convalescent plasma/serum were diluted in serum-free media for a final dilution of 1:40 and added in triplicate to a white-bottomed 96-well plate and titrated threefold. Serial dilution of monoclonal antibody (mAb) were prepared. Convalescent plasma/serum or mAbs were incubated with a fixed titred volume of sarbecovirus pseudoparticles equivalent to 10^5^ – 10^6^ signal luciferase units and incubated for 1h at 37°C and 5% CO_2_. Target HEK293T cells expressing human ACE2, or the relevant bat ACE2 receptor (*Rh. alcyone* for RhGB07 and RfGB02, *Rh. cornutus* for Rc-o319) were then added (at a density of 2×10^5^cells/mL for convalescent serum/plasma, and at 4×10^5^cell/mL for mAbs) and incubated at 37°C, 5% CO_2_ for 72 h. Firefly luciferase activity was then measured after the addition of Bright-Glo luciferase reagent on a luminescence plate reader (GloMax-Multi+ Detection System (Promega) for convalescent serum/plasma or Victor X3 multilabel plate reader (PerkinElmer) for mAbs). Pseudotyped virus neutralisation titres were calculating by interpolating the point at which there was a 50% reduction in luciferase activity, relative to untreated controls (IC_50_).

### Cell–cell fusion assays

HEK293T rLuc-GFP 1–7 [28] effector cells were transfected in OptiMEM (ThermoFisher Scientific) using Transit-X2 transfection reagent (Mirus Bio), as per the manufacturer’s recommendations, with sarbecovirus Spikes (500ng) alongside mock-transfection with empty plasmid vector (pcDNA3.1+) (**S1 Table**). BHK-21 rLuc-GFP 8-11 target cells were transfected with 500 ng of different ACE2-expressing constructs (**S2 Table**). Effector and target cells were co-cultured at a ratio of 1:1 in white 96-well plates to a final density of 4 × 10^4^ cells/well 2 days post transfection. Quantification of cell–cell fusion was measured based on Renilla luciferase activity, 24 h later by adding 1 μM of Coelenterazine-H (Promega) at 1:400 dilution in PBS. The plate was incubated in the dark for 2 min and then read on a Glomax Multi+ Detection System (Promega) as above and CSV files were exported for analysis.

### Recombinant protein production

Secreted sarbecovirus RBDs and mammalian ACE2 proteins were produced by cloning into the pOPIN-BAP-His and pOPIN-Fc expression vectors, respectively. These plasmids were then expressed in Expi293F cells using the PEI40K transfection reagent (Polysciences) with additives (300mM valproic acid, 500mM of sodium propionate, 2.5M glucose). Supernatants were harvested 4 days post-transfection, centrifuged for 45 mins at 6000 x g and filter through a 0.45µM filter. Presence of secreted protein was determined by Coomassie and by blotting for anti-His or anti-Fc. Supernatants were purified using HisTrapHP or HiTrap Protein A columns (Cytivia), desalted using Zeba Spin Desalting Columns (ThermoFisher Scientific) and concentrated using Amicon Ultra 15 columns (Merck Millipore, #C7715). The proteins were then quantified using a Nanodrop and a Pierce BCA Protein assay (ThermoFisher Scientific) and run on an SDS-PAGE gel to assess protein purity by Coomassie.

### Structural analysis

#### Protein Production

For protein production, this was performed as previously reported for Spike and ACE2 [10]. Briefly for Spike, a CHO-codon-optimized sequence of the soluble ectodomain of RhGB07 (residues 1 – 1191) was placed downstream of the u-phosphatase signal secretion peptide. Following the Spike sequence, a C-terminal T4-foldon trimerization domain, a 3C-protease cleavage site, 8x-His tag and a 2x Strep tag were included. A putative polybasic site RAK (residues 669-671) was mutated to GAS. A prefusion state 2P-stabilizing mutation was also added (residues 969-970). The protein was produced ExpiCHO cells and purified via the 2x Strep tag. For ACE2 expression and purification were also performed in the same manner as Spike.

### Cryo-electron microscopy

Cryo-EM grids were prepared as previously (PLOS PATHOGENS) on a Vitrobot Mark IV (Thermofisher Scientific (TFS)). Quantifoil R1.2/1.3 Au 400 holey carbon grids were glow-discharged for 90 s at 15mA using a PELCO easiGlow device (Ted Pella, Inc.). The putative complex was formed by mixing 3 uM of RhGB07 spike with 6 uM of either human or *Rh. ferrumequinum* ACE2 and applying 3 uL to the glow-discharged grids blotting for 6 seconds, blot force 10, at 95% humidty and 10C in the chamber. The blotted grid was plunge frozen in liquid ethane cooled by liquid nitrogen.

#### Data screening, collection and processing

On a TFS Titan Krios G4, the grids were screened and only one was suitable for data collection containing the putative RhGB07 Spike with human ACE2. On-the-fly processing on cryoSPARC live v3.3.2 was used during data acquisition (10.1038/nmeth.4169). Pre-processed motion corrected micrographs were patch corrected and used thereafter. 5 254 208 particles were template picked using PDB : 7QO7 as a template. After 2 rounds of 2D-classification clear classes showing a trimeric spike were observed comprising 2 685 particles. These particles were sufficient to produce an ab-initio model and can be refined to 3.66 Å with C3 symmetry. To increase the resolution and quality of the map these 2D classes were used once again for template picking which led to 2 234 763 particles. After several rounds of extracting, cropping and classification only 21 821 particles remained (3.09 Å) and were subjected to heterogenous refinement with 4 classes. The 3 best classes in terms of resolution were used (17 685 particles) for a final round of NU-Refinement leading to a map of 2.97 Å (FSC 0.143, C3 symmetry). This map showed good local resolution at the RBD without necessitating for local refinement for this domain.

#### Model building

The final model was built by docking Alphafold2 (10.1038/s41586-021-03819-2) predictions as individual domains (NTD, RBD, S2) using phenix.voyager (10.1107/S2059798323001602) followed by cycles of manual model-building in COOT (10.1107/S0907444910007493), specifically within the NTD, RBD and the putative basic region of the RhGB07. Analysis and figures were done within ChimeraX (10.1002/pro.4792).

#### Surface Plasmon Resonance

Kinetics and epitope binning assays were performed on the Carterra LSA^XT^ using HC30M chips. Chips were pre-conditioned using sequential 60s injections of 50 mM NaOH, 1M NaCl, and 10 mM pH2 glycine. For the kinetics assay, an anti-human Fc lawn was prepared by covalently coupling goat anti-human-Fc polyclonal antibody (BioRad 5211-8004) to the chip surface. Free carboxyl groups on the chip surface were activated for 5 minutes using 1-ethyl-3-(3-dimethylaminopropyl)carbodiimide hydrochloride (EDC) and 115 mg N-hydroxysuccinimide (NHS) (final concentrations of 130 mM and 33 mM respectively in PBS). Goat anti-human-Fc was diluted to 50 μg/ml in 10 mM sodium acetate (NaOAc) to pre-concentrate the protein into the negatively charged mix and applied to the chip for 7 minutes in two independent locations. The coupling reaction was quenched for 5 minutes using 500 mM ethanolamine, and the chip washed with three 30 second pulses of pH 1.9 glycine (10 mM). The human anti-RBD monoclonal antibodies were diluted in PBST (PBS + 0.005 % tween 20) to 1 ug/ml and 0.25 ug/ml and captured to the anti-human-Fc lawn for 7 minutes. Non-regenerative kinetics was performed using RBD diluted two-fold in PBST (5 μg /ml to 2.4 ng/ml) allowing a seven-minute association and 14-minute dissociation. After the highest concentration of RBD the anti-human Fc lawn was regenerated using three 30 second injections of pH 1.9 glycine (10 mM) and fresh human anti-RBD antibodies were captured for each sarbecovirus RBD kinetics cycle. Kinetic analysis was performed in Kinetics^TM^ v1.9.2 to calculate K_D_ by fitting to the 1:1 model following channel and double referencing. Average K_D_ was calculated using the values obtained from both coupling concentrations and the two independent couplings on the chip giving n = 4.

For epitope binning, human anti-RBD monoclonal antibodies were diluted to 1 μg/ml in NaOAc and directly coupled to the pre-conditioned HC30M chip as above allowing a 10 minute coupling period. SARS-CoV-2 wild type RBD was diluted to 0.625 μg/ml in PBST and injected for 5 minutes, followed by the competing antibody at 1 μg/ml for 5 minutes. The chip was regenerated using pH 1.9 glycine (10 mM) as previously before injecting the next RBD and competitor antibody. Epitope binning and community analysis was performed in Epitope^TM^ v1.9.2.

### Flow cytometry

BHK-21 cells were transfected using Transit-X2 transfection reagent (Mirus Bio), as per the manufacturer’s instructions with 500 ng of each ACE2-expressing construct (**S2 Table**) or mock-transfected with empty plasmid vector (pDISPLAY) for 48 h. Cells were resuspended in cold PBS and washed in cold stain buffer (PBS with 1% BSA (Sigma-Aldrich), 0.01% NaN_3_ and protease inhibitors (ThermoFisher Scientific). Cells were stained with anti-HA Phycoerythrin (PE)-conjugated antibody (Miltenyi Biotech) at 1:500 dilution for 10^6^ cells for 30 min on ice. In parallel, 10ug RBD recombinant proteins were stained with anti-His-Allophycocyanin (APC)-conjugated antibody (Miltenyi Biotech) at 1:500, and 50ul of stained RBD were mixed with cells for 1h to allow binding. Cells were then fixed with 2% paraformaldehyde (Fisher Scientific) on ice for 20 mins, then resuspended in PBS before being analysed using the MACSQuant Analyzer 10 (Miltenyi Biotech), and RBD-ACE2 binding was statistically calculated by SE Dymax % Positive. The mean SE Dymax % Positive was calculated from three biological replicates and analysed using FlowJo_v10.8.1. The gating strategy can be found in **Supp Fig 4A** and the same gating strategy was applied in all experiments.

### Western blotting

BHK-21 cells were transfected using Transit-X2 transfection reagent (Mirus Bio), as per the manufacturer’s instructions with 500 ng of different ACE2-expressing constructs (**S2 Table**) or mock-transfected with empty plasmid vector (pDISPLAY) and harvested 48 h post transfection. Pseudotyped virus was purified and concentrated under a 20% sucrose cushion by ultracentrifugation (23,000 rpm for 2h, 4°C). All protein samples were generated using 2× Laemmli buffer (Bio-Rad) and reduced at 95°C for 5 min. Samples were resolved on 7.5% acrylamide gels by SDS-PAGE, using semidry transfer onto nitrocellulose membrane. Blots were probed with anti-HA primary antibody (Cell Signalling, #2367) for ACE2-transfected cells at 1:1,000 or with anti-GAPDH (Cell Signalling, #2118) internal control at 1:3000. Purified psuedovirus was resolved using an anti-FLAG primary antibody (Cell Signalling, #2368) at 1:1,000, anti-Spike antibody (Novus Biologicals, #NB100-56578) at 1:500 or anti-p24 (ThermoFisher Scientific, #MA1-71515) as an internal lentiviral control at 1:1,000 in PBS-Tween 20 (PBS-T, 0.1%) with 5% (w/v) milk powder overnight at 4°C. Blots were washed in PBS-T and incubated with anti-mouse (for HA and p24) or anti-rabbit secondary (for GAPDH, Spike and FLAG), antibody conjugated with horseradish peroxidase (Cell Signalling, #7076 and #7074, respectively) at 1:3,000 in PBS-T for 1 h at room temperature. Membranes were exposed to Clarity Western ECL substrate (Bio-Rad) according to the manufacturer’s guidelines and exposed to autoradiographic film.

### RBD ELISA

Recombinant RBD was coated onto 96-well ELISA plates at 1µg/ml per well in carbonate/bicarbonate coating buffer (0.6M pH9.6) at 4-8 °C overnight. Plates were blocked for 1 at RT with PBS-T containing 1% BSA, after which a 3-fold dilution series of monoclonal antibodies (starting concentration 5µg/ml) or polyclonal COVID-19 plasma/serum (starting concentration 50µg/ml), or a 4-fold dilution series of ACE2 recombinant proteins (starting concentration 8µg/ml), were diluted in PBS and added for 1h, at 37°C. Plates were washed 3 times with PBS-T, followed by the addition of anti-human-Fc HRP conjugate diluted 1:10,000 added for 1h at 37°C. 1-step Ultra TMB (3,3’,5,5’-40 Tetramethylbenzidine) (Thermo Fisher Scientific) was added to each well, incubated for 5-10 mins at RT and the reactions stopped with an equivalent volume of 1M Sulphuric acid solution, after which the optical density (OD) at 450nm was measured using a GloMax Microplate Reader (Promega). Samples were analysed by plotting 50% effective concentration (EC50) using GraphPad Prism (8.2.1) software.

## Acknowledgements

COVID-19 convalescent plasma/serum samples were provided as part of the WHO collaborative study for the development of International Standard for anti-SARS-CoV-2 antibody coordinated by the MHRA (NIBSC) in partnership with CEPI. We also acknowledge The Pirbright Institute’s Flow Cytometry Facility and the Pirbright Institute Cell Servicing Unit. Additional thanks to Jonathan Popplewell at Carterra for his support with the set-up of the assay. We would like to thank Emiko Uchikawa, Alexander Myasnikov, Bertrand Becket and Sergey Nazarov from the Dubochef Center for Imaging (EPFL, UNIGE, UNIL initiative) for cryoEM grid preparation and data collection.

## Author contributions

**Conceptualisation:** DB, NT; **Formal analysis**: DB, NT, JN; **Funding acquisition**: DB, KJD, KL; **Investigation:** NT, JN, DN, JS, ALH, YL, AU, BEE, TPP, KL; **Methodology:** DB, NT, JN, JS, KJD, KL; **Project administration:** DB, NT; **Writing – original draft:** DB, NT; **Writing – review and editing:** all authors.

## Results

### Different bat sarbecovirus clades show varied bat ACE2 usage

Sarbecoviruses have been isolated from different bat species across Afro-Eurasia, phylogenetically clustering into established virus clades, when aligned using the receptor binding domain (RBD) amino acid sequences. Representative viruses from each clade were selected for analysis based on sequence availability and experimental data in publications [5, 17, 30, 31] (**Figure 1A**; highlighted in red). This included the clade Ia viruses SARS-CoV-1, WIV-1 and Rs4231 (yellow), clade Ib viruses SARS-CoV-2, RaTG13, BANAL-20-52, BANAL-20-236 and BANAL-20-103 (orange), clade II viruses RacCS203, RPYN06, Rf1/2004 and Rp3/2004 (blue), clade III viruses RhGB07, RfGB02 (green), and the clade V virus Rc-o319 (red). Clade IV viruses were not included in this study. The selected viruses have been isolated primarily in Asia (clade I, II, IV and V), and in parts of Europe and Africa (clade III), as indicated (**Figure 1B**). The bat species from which these viruses were isolated are globally widespread, (individual geographic distributions of each species are shaded as shown; overlay in **Figure 1B** and individually in **Supp Figure 1**). Of note, Rhinolophus species*, Pteropus giganteus, Lyroderma lyra, Rousettus leschenaultii and Taphozous melanaopogon* bat species overlap in their geographic reach, particularly in Southeast Asia, whereas the *Desmodus rotundus, Antrozous pallidus and Myotis lucifugus* species intersect in their range only in the Americas (**Figure 1B**).

**Figure 1:**
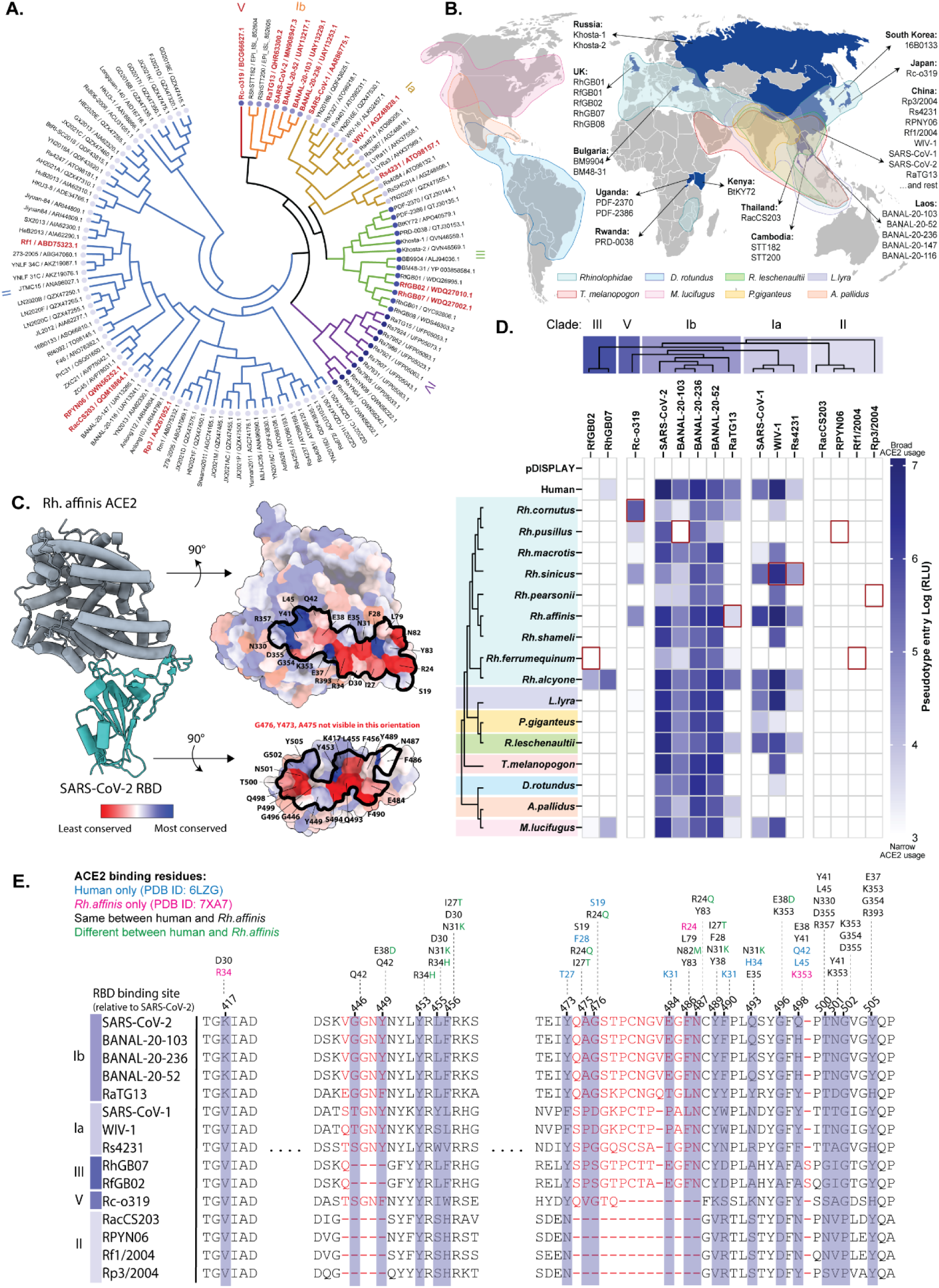
Phylogenetic and geographic distribution of bat sarbecoviruses and their bat ACE2 tropism. **(A)** Evolutionary history of bat sarbecovirus RBDs (NCBI reference shown for 110 amino acid sequences) was inferred by using the Maximum Likelihood method and Whelan and Goldman model. Initial tree(s) for the heuristic search were obtained automatically by Neighbour-Join and BioNJ algorithms to a matrix of pairwise distances (225 total positions) estimated using the JTT model with discrete Gamma distribution (5 categories (+G, parameter = 0.3460)), and then selecting the topology with superior log likelihood value. Evolutionary analyses were conducted in MEGA 11. Of note, the HKU3-7 and HKU3-8, and the HKU3-1, 2, 3, 4, 5, 6, 7, 9, 19, 11 and 12 Spike sequences are identical, so the HKU3-8 and HKU3-1 sequences were used as representatives, respectively. Bat sarbecovirus RBDs cluster into clades: clade Ia (yellow), clade Ib (orange), clade II (blue), clade III (green), clade IV (purple) and clade V (red). Viruses selected for analysis in this study are highlighted in red. **(B)** World map highlighting the geographic location where sarbecoviruses investigated in this study were first isolated (coloured in blue), along with the distribution of the bat species these were isolated from (https://www.iucnredlist.org/). **(C)** Structural images showing the binding of *Rhinolophus affinis* ACE2 with SARS-CoV-2 RBD (PDB 7XA7). Offset images highlight the conserved amino acid residues across the different bat ACE2s relative to Rh. affinis ACE2, and the conservation of bat sarbecovirus RBDs compared to SARS-CoV-2, screened in this study. The ACE2-RBD interacting residues are denoted. **(D)** Maximum likelihood phylogeny of the bat sarbecoviruses full length Spike and bat ACE2 amino acid sequences are shown. Bat sarbecovirus Spikes were used to generate lentiviral-based pseudotypes and used to infect BHK-21 cells overexpressing different bat ACE2s or human ACE2. A heatmap illustrating receptor usage is shown, representing the mean log RLU of 3-5 separate experiments. A vector only control (pDISPLAY) was included to demonstrate specificity and to set the background signal. The cognate bat receptor for each virus, where possible is highlighted with a red box. Absence of a red square is indicative of the lack of cognate receptor from our screen, either due to unavailability of ACE2 sequence, or because the original bat host is yet unknown (RhGB07: *Rh. hipposideros*; SARS-CoV-2, SARS-CoV-1: unknown; BANAL-20-52: *Rh. malayanus*; BANAL-20-236: *Rh. marshalli*; RacCS203 – *Rh. acuminatus*). **(E)** Sequence alignment of bat sarbecovirus RBDs at the ACE2-binding interface, with key amino acid residues denoted, using SARS-CoV-2 numbering. Regions of amino acid deletions between sequences are denoted in red. The corresponding human and *Rh.affinis* ACE2 residues that bind SARS-CoV-2 RBD are highlighted, with black indicating amino acid residues that are the same between human ACE2 and *Rh. affinis* ACE2, green indicating the same residues but a different amino acid, and the remainder highlighting unique binding residues for human ACE2 (blue) or *Rh. affinis* ACE2 (pink) only.

To understand how bat ACE2 tropism varies across the sarbecovirus sub-genus, a receptor usage screen was conducted in cells overexpressing human ACE2 (hACE2) or 16 different bat ACE2 receptors, which included a range of *Rhinolophus* species (*Rh. cornutus, Rh. pusillus, Rh. macrotis, Rh. sinicus, Rh. pearsonii, Rh. affinis, Rh. shameli, Rh. ferrumequinum, Rh. Alcyone*) and representative bat species from other families (*L. lyra, P.giganteus, R. leschenaultii, T. melanopogon, D. rotundus, A. pallidus and M. lucifugus)*. When their sequence conservation was analysed in the context of the structure of SARS-CoV-2 RBD bound to *Rh. affinis* ACE2 (PDB 7XA7), the selected bat sarbecoviruses and bat ACE2s showed high levels of divergence at the binding interface, suggesting that the receptor usage of these viruses with bat ACE2s assessed in this study would likely vary (**Figure 1C)**. Transfected cells were subsequently transduced with bat sarbecovirus Spike-based pseudotypes (those highlighted in red in **Figure 1A, S1 and S2 Tables**). Expression of ACE2 proteins in transfected cells, as well as SARS-CoV-2 Spike and p24 (internal lentivirus control) in purified pseudotypes, was confirmed in parallel (**Supp Fig 2**). To summarise, a diverse pattern of receptor usage was seen across the viruses tested (**Figure 1D**). Spike proteins from clade Ib sarbecoviruses, which includes SARS-CoV-2, had a broader (more *generalist*) receptor usage pattern, with RaTG13 being a notable exception. Indeed, only a few species of bat ACE2s were less permissive to SARS-CoV-2 or BANAL Spike-mediated entry. Clade Ia, which includes SARS-CoV-1, had a more intermediate receptor usage pattern, with clearer examples of restriction, even within the Rhinolophus bat family, e.g. *Rh. ferrumequinum* and *Rh. pusillus*. Nevertheless, WIV-1 and Rs4231, which were both isolated from *Rh. sinicus*, showed robust usage of this ACE2 receptor. In contrast, the clade III and V Spikes exhibited more restricted (*specialist*) ACE2 usage, primarily using the cognate bat ACE2 from which they were isolated, e.g. *Rh. cornutus* ACE2 usage by Rc-o319. As expected, the clade II viruses were unable to use any of the bat ACE2 receptors tested (**Figure 1D**) [1, 2, 5, 18, 32, 33]. Sequence alignment of the bat sarbecovirus RBDs, relative to SARS-CoV-2, revealed various deletions: at positions 446-449 in clade III viruses, at positions 480-487 for clade V viruses, and at positions 445-449 and 475-487 for clade II viruses (equivalent SARS-CoV-2 Wu-hu-1 numbering used throughout this study). The RBD-ACE2 interaction sites of *Rh. affinis* and human ACE2 were also compared based on known structures (human ACE2: PDB 6LZG, *Rh. affinis* ACE2: PDB 7XA7), noting differences in amino acid residues at positions 24, 27, 31, 34, 38 and 82. We also noted unique binding sites for human ACE2 that were not present for *Rh. affinis* ACE2, and vice versa (**Figure 1E**).

**Figure 2:**
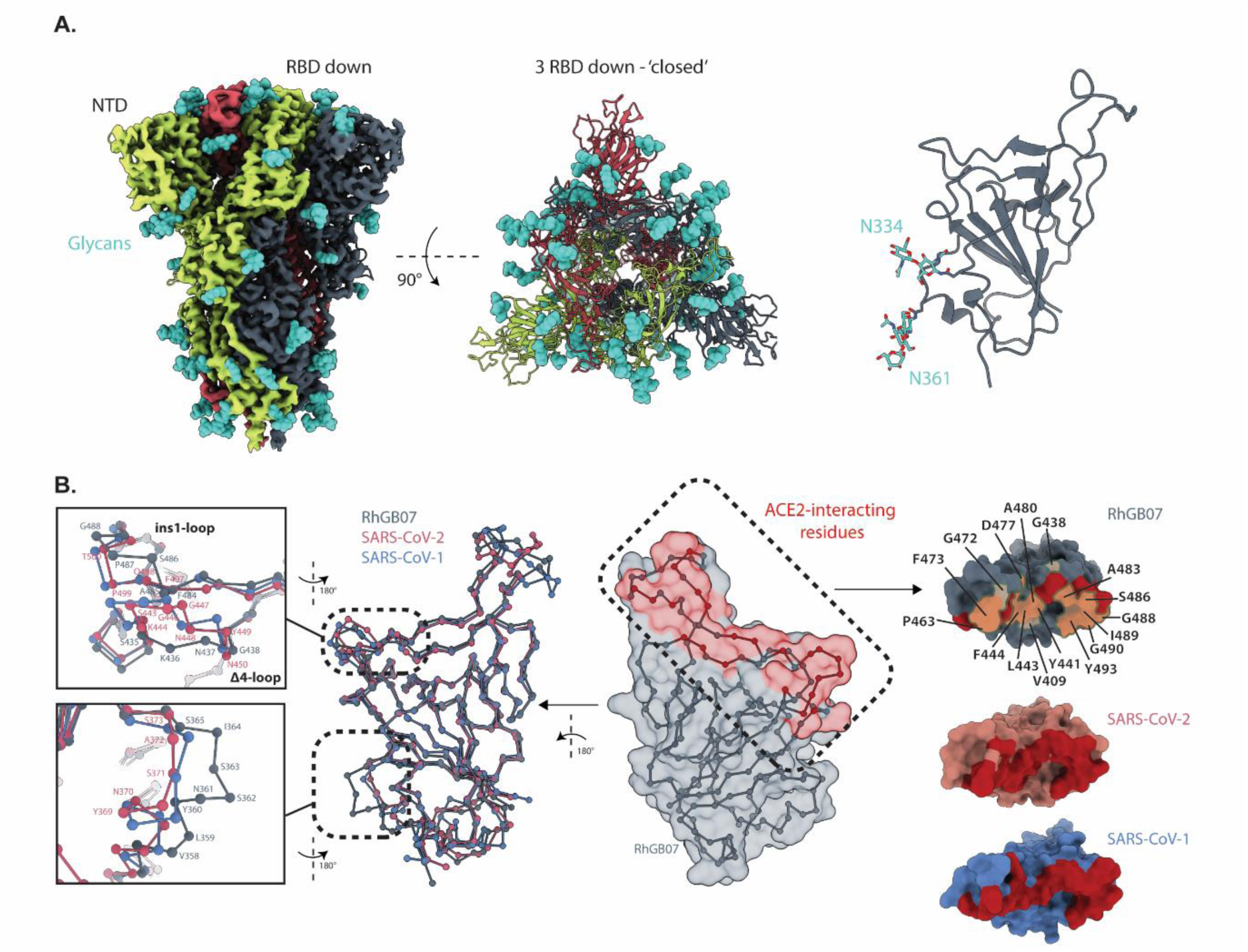
Cryo-EM structure of RhGB07 trimeric spike in a 3-RBD down state. **(A) left:** CryoEM density of the trimer spike coloured by the protomer chains the density originates from. Glycans are highlighted in teal. **Center:** the trimeric spike viewed from the top rendered in cartoon representation. **Right:** the isolated RhGB07 RBD represented as cartoon in grey with two glycans of interest shown. **(B)** The C-alpha trace of the RBD of RhGB07 rendered as balls-and-sticks superimposed with the RBD of SARS-CoV-1 and SARS-CoV-2. The insets show a zoomed-in view of two regions of the RBD that show the greatest changes in the position of the C-alpha of the RBD. The region containing the ins1-loop and delta4-loop coincides with putative ACE2-interacting residues based on mapping the equivalent residues of SARS-CoV-1 and SARS-CoV-2. The ACE2 interacting residues were determined by superimposing SARS-CoV-1 and RhGB07 RBD onto the SARS-CoV-2 RBD – ACE2 (PDB: 6M0J) structure and identifying interacting residues. Highlighted residues of RhGB07 show the putative interacting residues that differ in identity to SARS-CoV-2.

Coronavirus Spike proteins are also capable of inducing cell-cell fusion. To examine concordance between pseudotype entry and fusion, we repeated the assays in a quantitative cell-cell fusion assay, showing similar trends (**Supp Figure 3A**). Our cell-cell fusion assay also showed high concordance with the pseudotyped-based entry assay by Pearson’s correlation (**Supp Figure 3B**). The entry of bat sarbecoviruses with bat ACE2s also correlated to protein binding and affinity, i.e. those viruses that showed the highest levels of virus entry also bound most strongly in RBD-based flow cytometry assays (**Supp Figure 4A and B**). Further, purified recombinant bat sarbecovirus RBDs were used to assess binding to recombinant human ACE2 and two bat ACE2s (*Rh. cornutus, Rh. leschenaultti*) in ELISA; however these assays appeared to be less sensitive (**Supp Figure 4C**).

**Figure 3:**
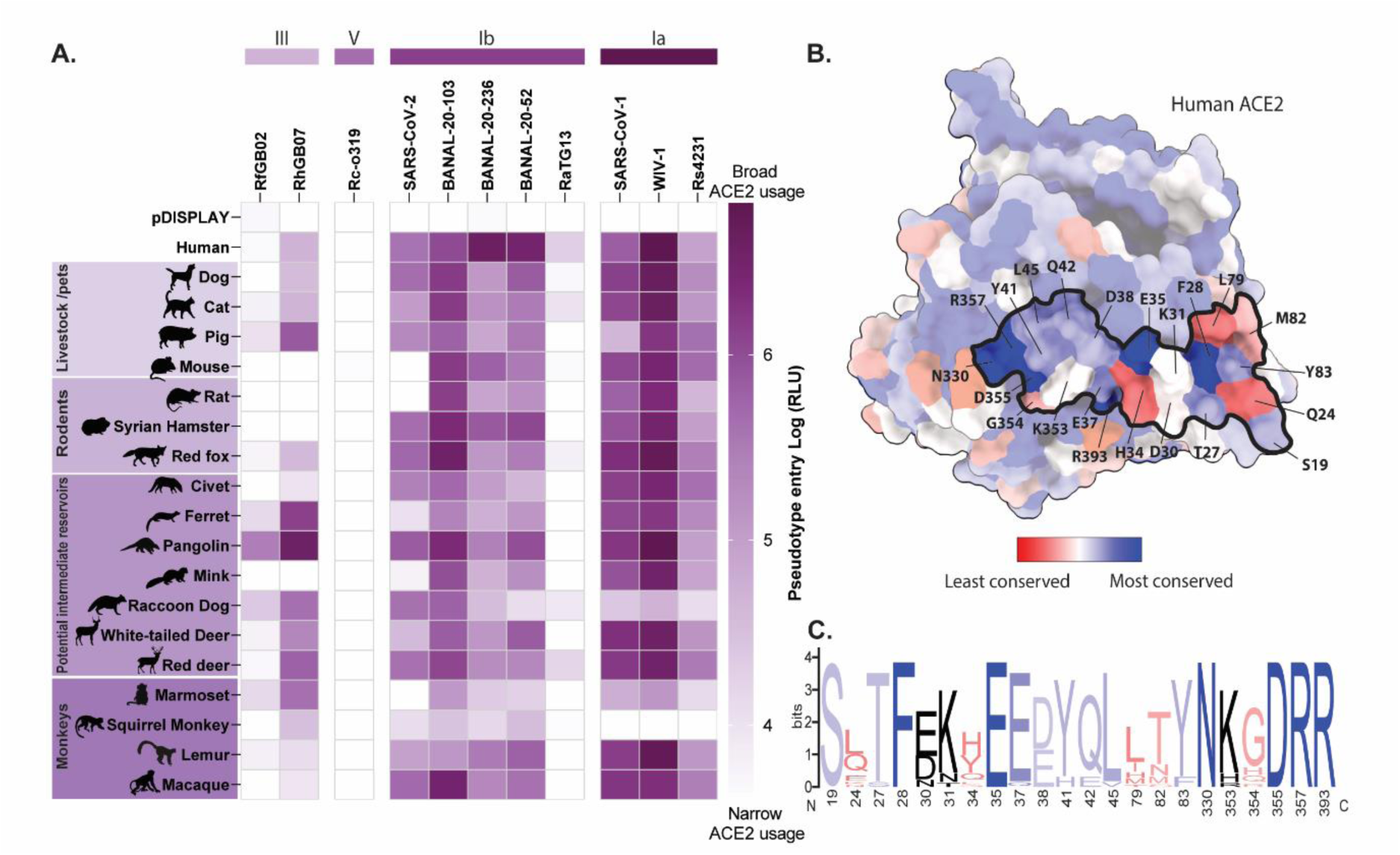
Bat sarbecovirus usage of mammalian ACE2. **(A)** Heatmap showing the receptor usage of bat sarbecovirus-pseudotyped viruses with mammalian ACE2s, including human, livestock/pets, rodents, potential intermediate hosts and monkeys. The mean log RLU from 3 separate experiments is shown and a pDISPLAY control to illustrate the background levels of entry. **(B)** Surface representation of human ACE2 coloured with mammalian ACE2 conservation (human ACE2: PDB 6M0J), with the outlined region denoting the RBD-interacting sites, with individual amino acids labelled. **(C)** WebLogo (University of California, Berkley, USA) plots summarising the amino acid divergence within the mammalian ACE2 sequences used in this screen. The single letter amino acid code is used with the vertical height presenting the relative frequency of that amino acid at each position. The positions at which ACE2 interacts with SARS-CoV-2 RBD are shown, N-terminal to C-terminal, with different colours representing amino acid conservation between the mammalian ACE2s examined in this study.

**Figure 4:**
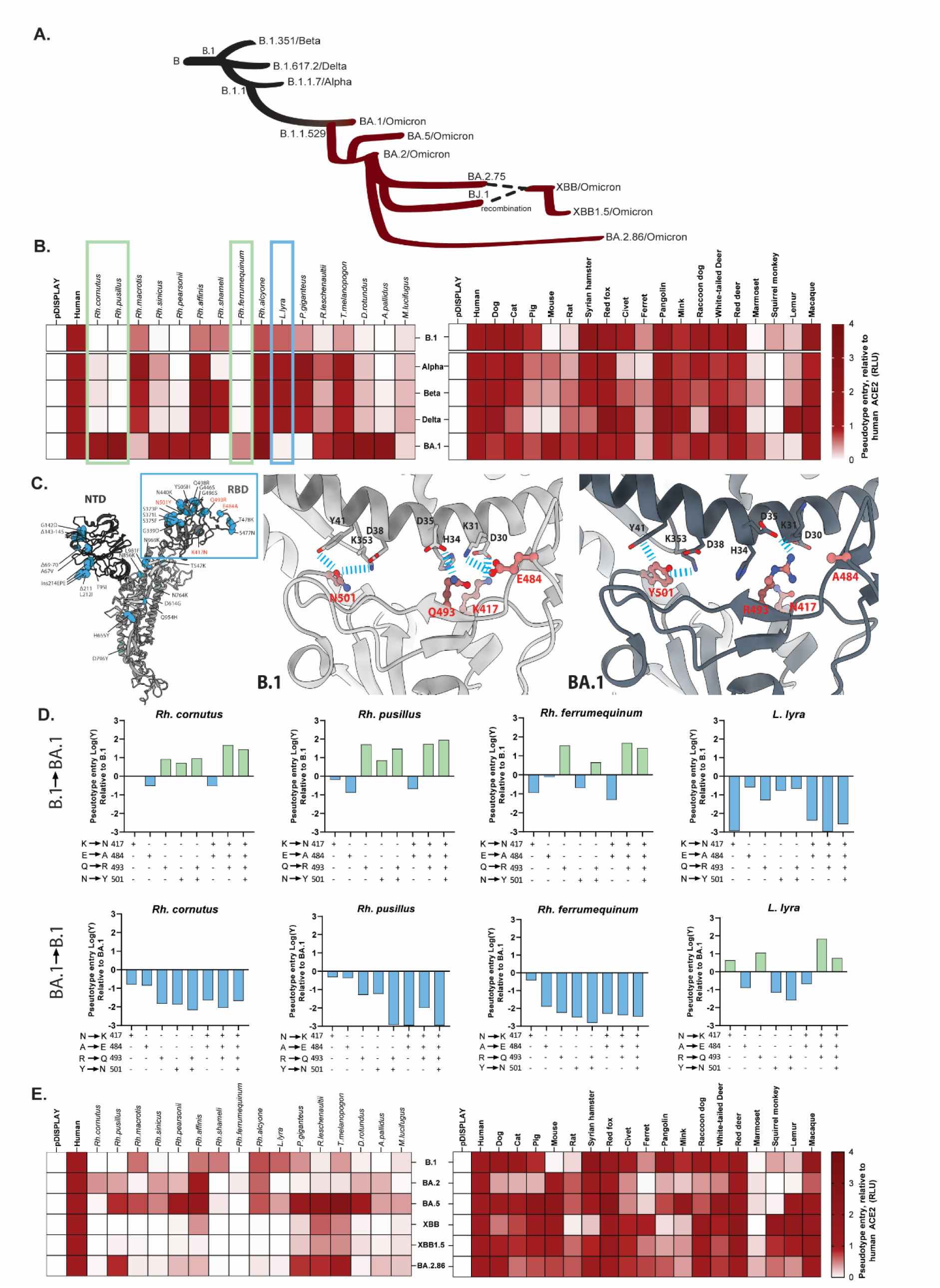
Species tropism of bat and mammalian ACE with SARS-CoV-2 variants. **(A)** Schematic of the emergence of SARS-CoV-2 variants depicting the divergence of the B lineage variants, including those variants of concern that grew to global dominance (B.1, Alpha, Beta, Delta, Omicron [black lines]), and the Omicron subvariants [red lines], some of which emerged via recombination events (dotted lines). Heatmap showing the bat and mammalian ACE2 usage of SARS-CoV-2 variants with **(B)** the B.1, Alpha, Beta, Delta and BA.1 variants, or **(E)** BA.2 and its subvariants BA.4/5, XBB, XBB1.5, BA.2.86. Data are shown as the mean fold change in pseudotype entry (RLU) of 3 separate experiments, relative to human ACE2. **(C)** Structure of SARS-CoV-2 BA.1 (PDB 7XO7) with mutations listed compared to B.1. Zoomed images of the Spike protein for B.1 (PDB 6M0J) and BA.1 (PDB 7WBP), highlighting key residues at positions 417, 484, 493 and 501 and the resultant interactions with human ACE2. **(D)** Specific amino acids within the B.1 RBD were mutagenised to make the RBD more BA.1-like (K417N, E484A, Q493R or N501Y) (top panel), or within BA.1 RBD to make the RBD more B.1-like (N417K, A484E, R493Q or Y501N) (bottom panel), either by themselves or in combination. Entry of these mutants was tested with *Rh. cornutus, Rh. pusillus*, *Rh. ferrumequinum*, and *L. Lyra* ACE2s. Data is shown as log fold-change in pseudotype entry, with green bars indicating an increase in virus entry and blue bars a decrease in entry, relative to B.1 (top panel) or BA.1 (bottom panel), highlighted also in Figure 4B.

### RhGB07 structure reveals a modified putative RBD-ACE2 interaction interface

Our functional screens revealed interesting differences between viruses within the same clade, particularly clade III viruses. For example, although RhGB07 exhibited a specialist receptor usage profile, it was able to enter cells overexpressing human ACE2, whereas RfGB02 could not (**Figure 1D**). To understand the structural determinants of generalism and specialism, as well as zoonotic potential across sarbecoviruses we therefore focused on resolving the structure of the specialist Spike, RhGB07. Previously, we produced recombinant 2P RhGB07 Spike with a putative basic region mutated and tested in BLI binding assays [10]. To reveal the molecular details of potential interactions between ACE2 and RhGB07 Spike, we attempted to determine a cryoEM structure of human ACE2 and RhGB07 Spike. We prepared grids for cryoEM by co-incubating either human or bat ACE2 (*Rh. ferrumequinum*) with RhGB07 Spike before freezing. Initial representative 2D classes did not reveal the presence of any bound ACE2 to Spike (**Supp Figure 5A, S3 Table**). Further classification revealed the presence of only 3-RBD closed/down Spike particles (in the dataset containing hACE2 and RhGB07). We could clearly define a limited number of particles and when C3 symmetry was imposed we could refine a map to 2.94Å (**Supp Figure 5A**). The model was built by docking Alphafold2 predictions as individual domains using phenix.voyager followed by manual model-building, specifically within the NTD, RBD and the putative basic region of the RhGB07. The structure revealed a trimeric organization of protomers, following a domain organization typical of sarbecoviruses. As noted in the 2D classes, the structure revealed a completely closed structure that shows no variation between RBD up or down (**Figure 2A**). Comparison with a recently described Clade III Spike, PRD-0038, that was also visualized as a 3-RBD down full-length Spike, shows a RMSD of 1.9 Å across all pairs of atoms suggesting high structural similarity between the two (**Supp Figure 6).** The RBD was less identical between PRD-0038 and RhGB07 with an RMSD of 2.1 Å. We note that we could also identify a similar number of glycans on the surface of RhGB07, in particular N361 (N360, PRD0038) that has been suggested as stabilizing a closed Spike conformation, as we see in our structure. Comparisons between the structures of the generalist sarbecovirus Spikes from SARS-CoV-1 and SARS-CoV-2 with RhGb07 revealed structural similarity except at the sites of deletion and insertions (**Figure 2B**). Within the RBD, we determined the loops that mediate ACE2 interaction in SARS-CoV-1 and SARS-CoV-2 adopt strikingly different 3D conformations as compared to RhGB07 and also contain residues of different identities to RhGB07 despite the overall RBD structure appearing highly identical (**Figure 2B**). For example, the ‘ins1-loop’ where there is a single residue insertion in RhGB07 compared to SARS-CoV-2, positions serine 486 (RhGB07 numbering) in place of the Q498 residue in SARS-CoV-2. Changes at these hotspot loops may be determinants for RhGB07-ACE2 binding selectivity.

**Figure 5:**
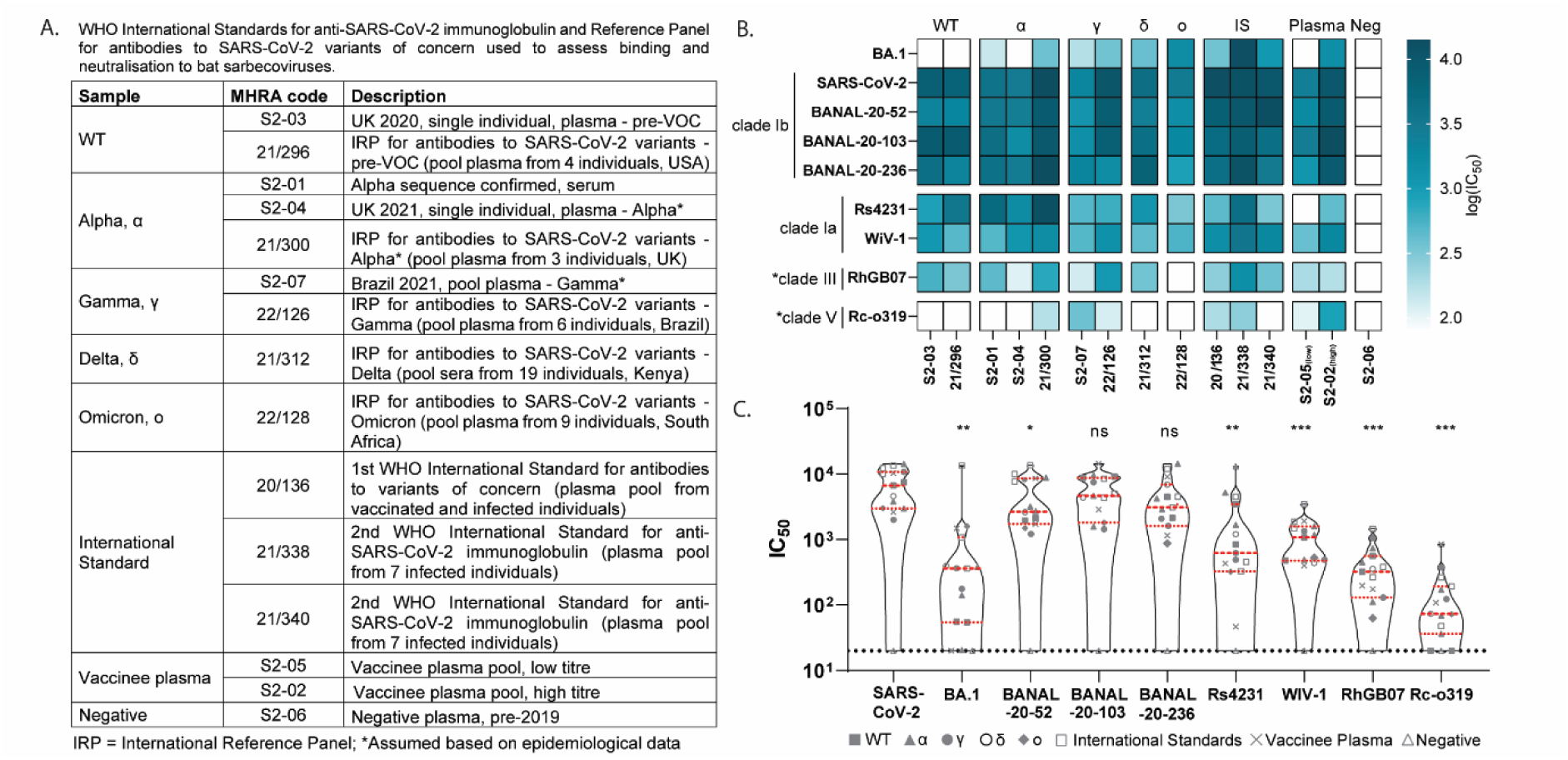
Antigenicity of bat sarbecoviruses using SARS-CoV-2 polyclonal convalescent sera. **(A)** Table outlining the different convalescent plasma/serum used to investigate the antigenicity of bat sarbecoviruses. These include samples from individuals following infection with the wildtype (WT), Alpha, Gamma, Delta, or Omicron BA.1 variants, either confirmed by PCR or assumed based on epidemiological data. The sample set also includes the WHO International Standards provided by MRHA (NIBSC), pooled vaccinee plasma (low and high titre) and negative plasma, pooled from individuals pre-2019. Information about the COVID-19 standards and reagents can be found here: [WHO/BS/2022.2427: Establishment of the 2nd WHO International Standard for anti-SARS-CoV-2 immunoglobulin and Reference Panel for antibodies to SARS-CoV-2 variants of concern](https://www.who.int/publications/m/item/who-bs-2022.2427). **(B)** Heatmap showing the log (IC_50_) values of neutralisation (the dilution at which there was a 50% reduction in luciferase activity) of SARS-CoV-2, BANAL-20-52, BANAL-20-103, BANAL-20-236, Rs4231, WIV-1, RhGB07, Rc-o319 and BA.1 viruses in the presence of the MHRA (NIBSC) reference reagents. HEK293T cells stably expressing human ACE2 were used for all neutralisation assays, apart from RhGB07 and Rc-o319, where HEK293T cells overexpressing *Rh. alcyone* or *Rh. cornutus* were used, respectively (indicated by an asterisk). **(C)** Violin plots show collated IC_50_ values for the neutralisation of these viruses with each sera set, with the median, upper and lower quartiles (red lines) and limit of detection (black dotted line) shown. A one-way ANOVA with Dunnet’s multiple comparison was performed, comparing to SARS-CoV-2, with significance values indicated (* = p<0.05, ** = p<0.005, ***= p<0.0005, ns = non-significant).

**Figure 6:**
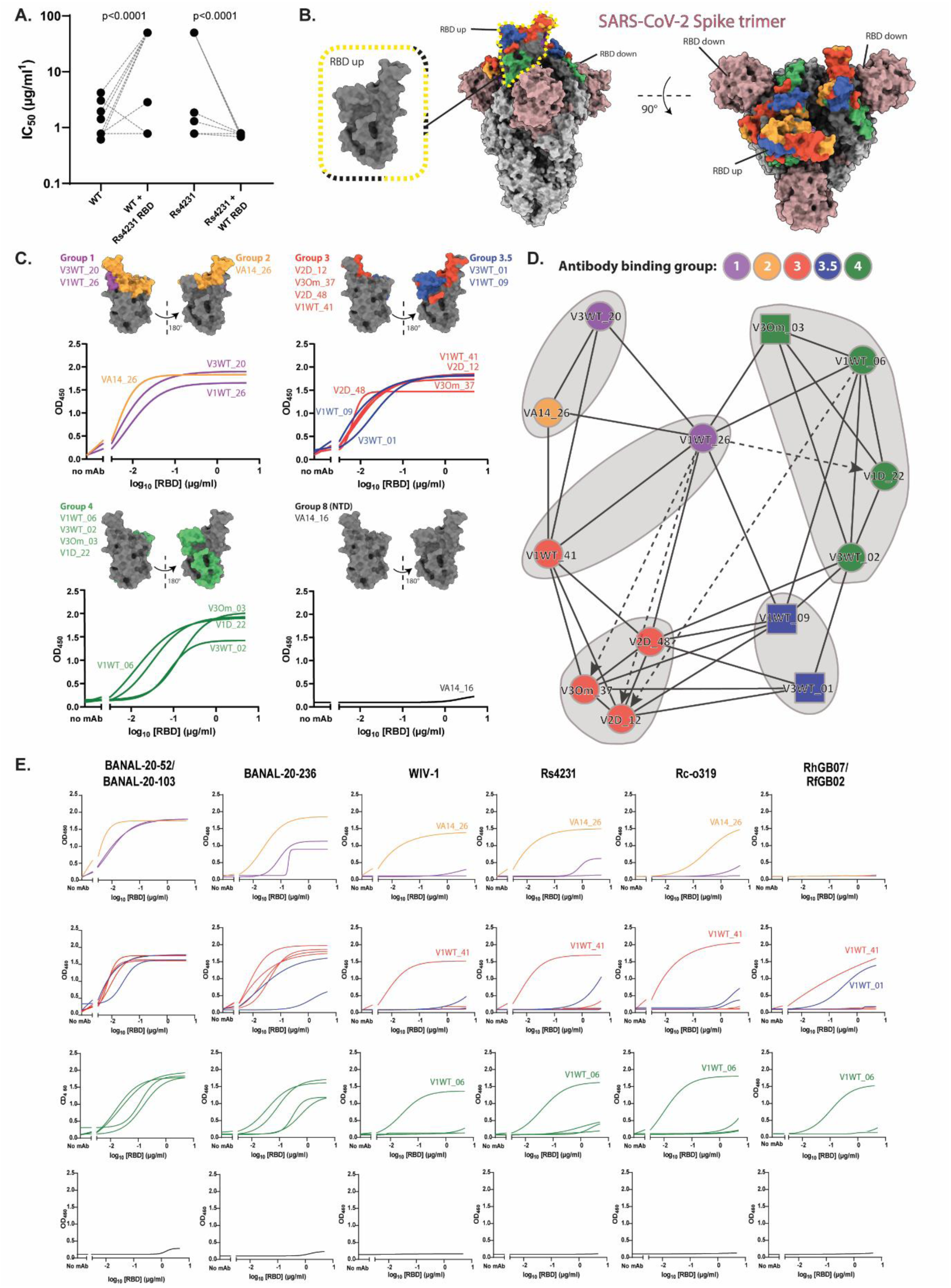
Binding of monoclonal antibodies isolated from SARS-CoV-2 breakthrough infections to bat sarbecovirus RBDs. **(A)** Neutralisation of SARS-CoV-2 and Rs4231 RBD chimeras with SARS-CoV-2 RBD-specific monoclonal antibodies (mAb) (IC_50_ µg/ml), with dotted lines linking matched samples. The p values are shown, calculated by comparing the change in mean IC_50_ value and by using Tukey’s multiple comparison tests, respectively**. (B)** Spike structure of SARS-CoV-2 with 2 RBD-down and 1 RBD-up (PDB 7QTI) with the binding epitope regions of the different mAb competition groups is shown with both a side- and top-view (Group 1 = purple, Group 2 = orange, Group 3 = red, Group 3.5 = blue, Group 4 = green, Group 8 = non-RBD, NTD-specific, defined previously [39, 51, 52]. **(C)** SARS-CoV-2 RBD structures (PDB 7QTI) highlighting a broad epitope footprint of the mAb competition groups corresponding binding data. Each of the plots shows a single mAb binding in an ELISA using purified **(C)** SARS-CoV-2 or **(E)** other bat sarbecovirus RBDs. Plots show raw optical density (OD) values with curves calculated using a least squares regression non-linear fit. **(D)** Epitope binning by surface plasmon resonance (SPR) identifies 5 epitope communities broadly aligning with the antibody competition groups. Grey circles indicate communities by setting the associated dendrogram threshold to 0.45. Solid lines indicate symmetrical blocking, regardless of which mAb is the ligand. Dotted lines indicate asymmetrical blocking, with the arrowhead indicating the direction of blocking, pointing towards the antibody immobilised as a ligand. Circles indicate antibodies for which ligand and analyte data was available, squares indicate only analyte data was available.

Together, these functional and structural data suggest that sarbecoviruses have distinct receptor usage profiles, differences which can likely be attributed to variations within the RBD-ACE2 interaction interface. To understand whether patterns of generalism or specialism in the bat reservoir host are conserved in potential intermediate reservoirs we expanded our analysis to a wider range of mammalian ACE2 proteins.

### Bat sarbecoviruses also have broad tropism for mammalian ACE2

Based on previously established ACE2 restrictions [15, 17, 18, 21, 34], evidence for a role in spillover of sarbecoviruses into humans [35–37], and/or the identification of SARS-CoV-2 reverse zoonotic spillover [25–27, 37, 38] we selected 19 additional mammalian ACE2s for analysis including human, livestock/pets (dog, cat, pig), rodents (mouse, rat, Syrian hamster), potential intermediate reservoirs (red fox, civet, ferret, pangolin, mink, raccoon dog, white-tailed deer, red deer), and monkeys (marmoset, squirrel monkey, lemur, macaque) (**S2 Table**). Clade II Spikes that did not use any of the screened bat ACE2s in the initial screen were excluded from further analysis. Similar to the bat ACE2s, clade Ib Spike pseudotyped viruses exhibited broad tropism with the majority of mammalian ACE2s tested, with notable restrictions for SARS-CoV-2 with mouse, rat, ferret, mink, marmoset and squirrel monkey ACE2, as published previously [15]. Interestingly, these restrictions were less obvious for the closely related BANAL viruses, including BANAL-20-52 whose RBD is almost identical to SARS-CoV-2, whereas RaTG13 was much more restricted. The mammalian ACE2 usage profile of the clade III specialists RfGB02 and RhGB07, as well as the clade V specialist Rc-o319, were more divergent than with the bat ACE2s. For example, whilst RhGB07 Spike was able to use the ACE2 from many mammals, including pangolin and raccoon dog to enter cells, Rc-o319 was entirely restricted with this panel. The lack of an absolute correlation between generalism and specialism in bats and other mammals was also seen for clade Ia pseudoviruses, which demonstrated almost universal usage of all ACE2s in our mammalian ACE2 receptor usage, aside from New World monkeys (**Figure 3A**). Comparison of mammalian ACE2 protein sequences (relative to human ACE2) revealed a much higher sequence conservation than seen for bat ACE2s (**Figure 1C**), particularly at the RBD-binding interface, with divergence observed at amino acid residues 24, 34, 79 and 82, and to a lesser extent at positions 30, 31 and 353 (**Figure 3B and C**).

### Bat and mammalian ACE2 usage has changed in line with continued SARS-CoV-2 evolution in humans

The D614G-containing SARS-CoV-2 lineage, B.1 was responsible for the first wave of the COVID-19 pandemic, with subsequent waves caused by genetically divergent variants including Alpha, Beta, Delta, and Omicron. All these variants, in particular Omicron, were defined by the introduction of important mutations within the Spike protein (**Figure 4A, Supp Figure 7A**). To examine changes to bat and mammalian ACE2 receptor usage by these SARS-CoV-2 variants we measured the entry efficiency of Spike-based pseudotypes in cells overexpressing ACE2 orthologues. For bat ACE2s, we observed enhanced (compared to B.1) bat ACE2 usage with the Alpha, Beta and Delta variants approaching or exceeding the levels recorded for human ACE2, however, the overall pattern of receptor usage did not change. In contrast, a divergent pattern was seen with Omicron BA.1, including shifted receptor usage of *Rh. cornutus, Rh. pusillus, Rh. ferrumequinum* and *L. lyra* ACE2 (**Figure 4B, left panel**).

**Figure 7:**
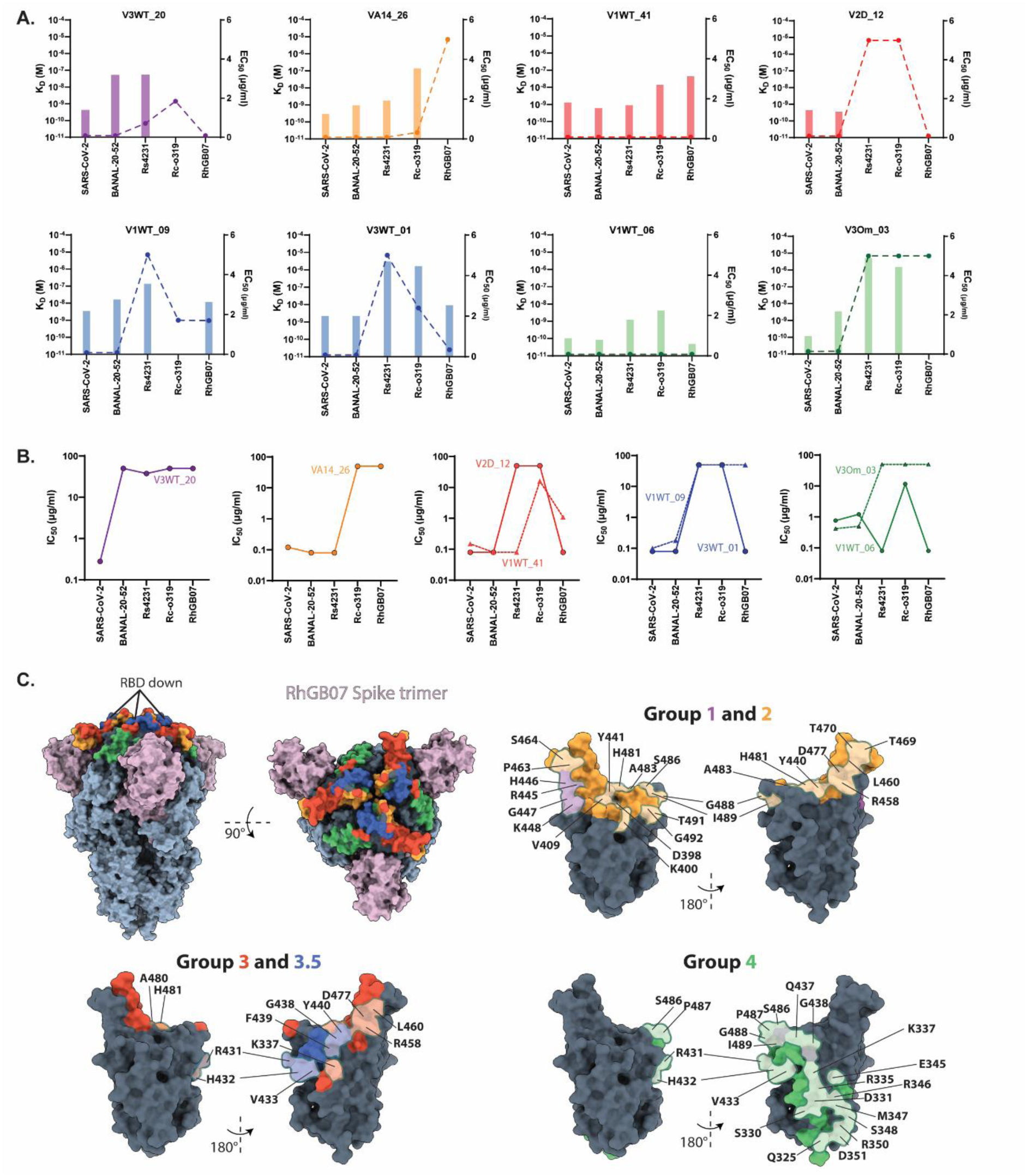
Antigenicity of bat sarbecoviruses using monoclonal antibodies isolated from SARS-CoV-2 breakthrough infections. **(A)** Dissociation constant (K_D_ (M), 2-4 replicates ± sd) of mAb binding kinetics calculated by SPR (bars) and ELISA EC_50_ (µg/ml, 2 replicates ± sd) values (broken lines with filled symbols) for a subset of mAbs and sarbecovirus RBDs. For K_D_, no bar indicates no binding by SPR. For EC_50_, lower limit of detection: <0.08µg/ml, upper limit of detection: >5µg/ml. **(B)** Neutralisation (IC_50_) titres shown for each mAb against bat sarbecovirus pseudotyped viruses (µg/ml), with limits of detection indicated (lower limit: <0.08µg/ml; upper limit: >50µg/ml for IC_50_). HEK293T cells stably expressing human ACE2 were used for all neutralisation assays. apart from RhGB07 and Rc-o319, where HEK293T cells overexpressing *Rh. alcyone* or *Rh. cornutus* were used, respectively. **(C)** RhGB07 trimer and RBD, with antibody competition groups highlighted, and are coloured with the same colouring as the SARS-CoV-2 trimer in (Figure 6B). Regions of divergence in binding epitopes in RhGB07 compared to SARS-CoV-2 are depicted, with amino acid residues denotes. Colours in all graphs/structures indicate mAb binding competition groups (Group 1 = purple, Group 2 = orange, Group 3 = red, Group 3.5 = blue, Group 4 = green). EC_50_, K_D_ and IC_50_ values be found in **S4 Table**.

In contrast, the species tropism for a wider selection of mammalian ACE2s was somewhat unchanged, maintaining the generalist properties of B.1, with the same notable restrictions, e.g. new world monkey (marmoset and squirrel monkey) and ferret. Notably, an increase in mouse and rat ACE2 usage was detected in all variants harbouring N501Y mutations in the Spike RBD, as previously shown (**Figure 4B, right panel**) [21].

Focusing on the significant differences in bat ACE2 receptor usage between B.1 and Omicron BA.1, we next investigated which mutations in the Spike were responsible. Compared to B.1, Omicron BA.1 has a significant number of amino acid substitutions in Spike, with 13 mutations in the RBD alone, particularly at residues that interact directly with ACE2, with the 417N and 484A mutations reducing potential interactions (**Figure 4C**). Based on our structural understanding of RBD residues that interact directly with ACE2, and/or previous evidence of a direct contribution to altered species tropism, we generated B.1→BA.1 substitution mutants (K417N, E484A, Q493R, N501Y) either individually or in combination. In addition, inverse mutations (BA.1→B.1) were built into a BA.1 backbone, with all mutants being functional when titrated on cells stably-expressing human ACE2 (**Supp Figure 7B**).

We then assessed the entry phenotype of these mutated Spikes with bat ACE2s that showed a switch towards increased (*Rh. cornutus, Rh. pusillus, Rh*. *ferrumequinum*) or decreased receptor usage (*L. lyra)* with BA.1 compared to B.1. We noted that whilst the K417N and E484A mutants on their own, or in combination, had little impact on receptor tropism, the Q493R and N501Y mutations, either in isolation or in combination with other mutations, resulted in increased receptor usage of *Rh. cornutus* and *Rh. pusillus* ACE2. For the *Rh. ferrumequinum* ACE2 the Q493R mutation alone was sufficient to switch receptor usage. For *L. lyra*, the decrease in receptor usage was largely attributable to the change at position 417 (**Figure 4D, top panel**). When the corresponding mutations were introduced into the BA.1 backbone, a similar phenotype was observed, with amino acid changes at positions 493 (R493Q) and 501 (Y501N) being detrimental to *Rh. cornutus Rh. pusillus* and *Rh. ferrumequinum* ACE2 receptor usage, especially when in combination. Likewise, the 417 (N → K) and 493 (R → Q) mutations in BA.1, as single mutations or together, were able to revert the observed restriction between BA.1 Spike and *L. lyra* ACE2 usage (**Figure 4D, bottom panel**).

Finally, we also examined the receptor usage of contemporaneous Omicron sub-lineages that have emerged and risen to dominance since 2022. Interestingly, aside from BA.4/5, and to a lesser extent BA.2.86, many of the Omicron variants tested (BA.2, XBB and XBB1.5), exhibited a reduced or complete loss of receptor usage for many bat ACE2 receptors (**Figure 4E, left panel**). This wasn’t evident for the wider library of mammalian ACE2 species, which remained widely used by the Omicron sub-linages tested (**Figure 4E, right panel**). Indeed, there was evidence of previous restrictions being overcome by variants such as XBB and BA.2.86, e.g. new world monkey and ferret ACE2s. In summary, the continued evolution of SARS-CoV-2 in humans has shifted the receptor usage pattern of this sarbecovirus away from an ancestral bat phenotype, towards one that is more generalistic for other mammals, including humans.

### Multiple bat sarbecoviruses share antigenic similarity to SARS-CoV-2

Several bat sarbecovirus Spikes in our study showed a broad host range for mammalian ACE2s, including human ACE2 (**Figure 3A**), suggesting the potential for spillover into human populations. To ascertain whether pre-existing immunity following infection with SARS-CoV-2 can provide cross-protection against related viruses we examined their antigenic similarity by ELISA and VNT with a panel of COVID-19 convalescent plasma/serum obtained from the Medicines and Healthcare products Regulatory Agency (MHRA), UK. The antibody panel includes the World Health Organization (WHO) International Standards for anti-SARS-CoV-2 immunoglobulins, and the International Reference Panel material for antibodies to SARS-CoV-2 variants of concern B.1 (wildtype, WT), Alpha, Gamma, Delta, and Omicron BA.1. We also tested plasma from vaccinees with high and low neutralising antibody titres to SARS-CoV-2, as well as negative plasma collected from healthy donors pre-2019 (**Figure 5A**). By ELISA we observed high levels of human IgG-based cross-specific binding to the RBDs of clade Ia and Ib viruses. Interestingly, we also noted binding to the RhGB07 and Rc-o319 RBD by most of the sera/plasma tested despite their phylogenetic divergence. (**Supp Figure 8**). The same sera/plasma also exhibited moderate-to-high neutralisation titres against equivalent pseudotyped viruses (BANAL-20-103, BANAL-20-236: ns; BANAL-20-52, * p<0.05; Rs4231: ** p<0.005; WIV-1: *** p<0.0005, compared to SARS-CoV-2) (**Figure 5B and C**). Due to the inefficient usage of human ACE2 by Rc-o319 and RhGB07 pseudotypes, the target cells for these virus neutralisation assays were instead transiently transfected with *Rh. cornutus* ACE2 and *Rh. alcyone* ACE2, respectively. Compared to SARS-CoV-2 we noted lower neutralisation titres against RhGB07 (***, p<0.0005), and lower still against Rc-o319 (***, p<0.0005) (**Figure 5B**, **Figure 5C**). Interestingly, the binding and neutralisation phenotype observed for these two viruses, which were the most antigenically distant in our screen, were similar to those observed for Omicron (BA.1). This is perhaps unsurprising given that most of the sera/plasma tested was taken before BA.1’s emergence; however, it does highlight the role of immune pressure in driving SARS-CoV-2 evolution (**Supp Figure 8A, Figure 5B**, **Figure 5C**). Another interesting observation was that in some ELISAs we did not detect binding e.g., S2-03 did not bind to the RhGB07 RBD, and, 21/296 and S2-04 did not bind to WIV-1 RBD, however we were still able to detect neutralisation of respective pseudotypes, suggesting neutralisation sites outside the RBD (**Figure 5B, Supp Figure 8**). In summary, both polyclonal sera and plasma from SARS-CoV-2 convalescent individuals shows robust evidence for the presence of cross-neutralising pan-sarbecovirus antibodies.

### Pan-sarbecovirus monoclonal antibodies are identifiable following breakthrough SARS-CoV-2 infection

Building on this cross-neutralisation data, we confirmed the RBD as the predominant target for this immune-recognition. Using SARS-CoV-2 as an exemplar ACE2 generalist and Rs4231 as a specialist (**Figure 1D**, **Figure 3A**, **Figure 5**) we generated RBD chimeras of these two Spikes and confirmed their functionality (**Supp Figure 9A**). We opted against the more phylogenetically and phenotypically distant viruses, RhGB07 and Rc-o319, because of their inefficient human ACE2 usage, which would limit direct comparison with SARS-CoV-2. Using a panel of RBD-specific monoclonal antibodies (mAbs) isolated from SARS-CoV-2 breakthrough infections [39], we then conducted neutralisation assays with these chimeras. We detected a reduction in IC_50_ when the Rs4231 RBD was introduced into the SARS-CoV-2 backbone (p<0.0001), whereas the converse was observed when the SARS-CoV-2 RBD was inserted into the Rs4231 backbone (p<0.001) (**Figure 6A**). We also confirmed that the differences in ACE2 receptor usage tropism between these two Spikes mapped to the RBD (**Supp Figure 9B**), e.g. switching mouse and marmoset ACE2 usage (**Supp Figure 9B**). These data confirmed the RBD as a major target for cross-specific antibodies, and that both receptor generalism/specialism and antigenicity functionally partition to the same Spike protein sub-domain.

The mAbs used in our screen have been characterised in more detail previously, based on binding and competition assays with mAbs to SARS-CoV-2 RBD (Group 1 [purple], Group 2 [orange], Group 3 [red], Group 3.5 [blue], Group 4 [green] and Group 8 [non-RBD binder]) [39] (**Figure 6B**). To confirm these mAb groups and to provide additional resolution, binding ELISAs and epitope binning competition assays were performed by assessing binding to SARS-CoV-2 RBD. In initial ELISAs, all RBD-specific mAbs bound to the SARS-CoV-2 RBD, whereas the NTD-specific mAb VA14_16 failed to recognise this protein, as expected (**Figure 6C**). Epitope binning was then performed by high throughput surface plasmon resonance (SPR) on the Carterra platform. In general, the same antibody binding groups were preserved, with additional blocking interactions between groups observed (**Figure 6D, Supp Figure 10**). Blocking interactions between group 3, 3.5, and 4 antibodies varied, indicating the presence of subtle differences in antibody binding within these groups (**Figure 6D**). For example, antibodies VA14_26 (group 2) and V1WT_41 (group 3) were characterised as having similar blocking phenotypes, indicative of epitope overlap (**Figure 6D**). Interestingly, the V1WT_26 antibody (group 1) showed blockage of group 2, 3, and 4 antibodies, despite group 1 mAbs occupying binding a distinct region of the RBD [39], suggesting possible steric hinderance (**Figure 6D**).

Building on the partial neutralisation of Rs4231 with these mAbs, we performed a more comprehensive assessment of their binding (by ELISA) phenotype, against other bat sarbecoviruses in our library. As with the polyclonal sera tested above, SARS-CoV-2, BANAL-20-52 and BANAL-20-236 RBDs were bound well by all the mAbs tested. In contrast, the more divergent WIV-1, Rs4231, Rc-o319 and RhGB07 RBDs were bound less robustly (**Figure 6E**). However, significant binding was maintained by antibodies in group 2 (VA14_26, except RhGB07), group 3 (V1WT_41) and group 4 (V1WT_06) (**Figure 6C**). Interestingly, these 3 mAbs were also able to bind the RBD of the non-ACE2 using clade II virus, RacCS203, albeit to a lesser degree (**Supp Figure 11**). As expected, none of the group 8 mAbs (NTD-specific) bound to any RBD (**Figure 6E**).

A representative subset of these sarbecovirus RBDs and mAbs were next analysed for complementary binding kinetics and affinity by ELISA and SPR, respectively. Generally, there was good correlation between the assays, wherein mAbs with low EC_50_ values also exhibited lower K_D_ (dissociation constant) values, indicated higher affinity (**Figure 7A**). However, some RBD/mAb interactions characterised as non-binding by ELISA showed low binding by SPR and a greatly increased K_D_ (**Figure 7A, S4 Table**). For example, V3Om_03 showed minimal to no binding to the more divergent RBDs in ELISA, but in SPR a low level of binding was detected for Rs4231 and Rc-o319 (**Figure 7A, S4 Table**). Interestingly, the converse trend was observed for mAb V3WT_20 with RhGB07 RBD, with binding identified by ELISA but not by SPR (**Figure 7A, S4 Table**).

Finally, we examined the neutralisation of various pseudotyped spikes by these mAbs. These experiments showed a similar trend, with BANAL-20-52 exhibiting binding EC_50_s and neutralisation IC_50_ values of <0.08 µg/ml, close to SARS-CoV-2 (**Figure 7B, S4 Table**). Mirroring the ELISA data, Rs4231 (clade Ia), RhGB07 (clade III) and Rc-o319 (clade V) were less well neutralised; however, the same three monoclonal antibodies (VA14_26, V1WT_41 and V1WT_06) showed good efficacy, highlighting competition groups 3, 3.5 and 4 antibodies as being important mediators of cross-neutralisation (**Figure 7B, S4 Table**). Neutralisation IC_50_ titres also correlated with K_D_, with larger a K_D_ correlating with weakened or abrogated neutralisation. In contrast, the three antibodies which exhibited broad RBD binding and neutralisation (VA14_26, V1WT_41, and V1WT_06) displayed a much narrower fluctuation in K_D_ consistent with their ability to bind all the RBDs tested (**Figures 7A and B, S4 Table**). Mapping the broad footprints of where these mAb competition groups might bind onto the Spike trimer of RhGB07 highlighted a significant number of amino acid residues that differ to SARS-CoV-2 RBD (**Figure 7C**). RhGB07 has a 3 RBD-down, closed conformation, which may contribute mAb accessibility, particularly the Group, 3.5 and 4 mAbs, which target the most exposed epitopes on in the RhGB07 RBD down/closed state (**Figure 7C**). Our companion article describes in more detail the structural basis for the broad specificity of these antibodies to sarbecovirus RBDs.

## Discussion

Identifying differences in receptor usage between closely related viruses and the key amino acid substitutions that underpin restriction are important tools for understanding the mechanisms of spillover. However, performing this work in a wholistic way, i.e. one that develops broad, translatable knowledge that improves our understanding of general rules governing spillover is more likely to have tangible benefits on pandemic prevention and preparedness. Herein, we describe that across the sarbecovirus sub-genus we could only find broad generalist ACE2-using viruses within the same clade (clade I) as SARS-CoV and SARS-CoV-2, and, that these viruses appear to all be antigenically similar. Interestingly, even for the more specialised ACE2-using sarbecoviruses we were still able to detect low levels of cross-neutralisation, which we could attribute to specific antibody classes. These data, combined with high post-pandemic levels of COVID-19 immunity in human populations, may mean the risk of future sarbecovirus zoonotic spillovers is diminished.

To arrive at this conclusion, we first assessed the pattern of generalism and specialism across the various extant clades of sarbecoviruses. Like SARS-CoV-2, other clade Ib viruses were mostly bat ACE2 generalists. In contrast, clade Ia and to a greater extent, clades III and V were more specialised, using primarily the ACE2 receptor of the bat species from which the virus was isolated. Interestingly, within clade Ib, RaTG13 showed a distinct, more restricted pattern ACE2 usage, despite sharing 98% amino acid similarity with SARS-CoV-2 and clustering in the same clade as SARS-CoV-2 and the BANAL viruses. This is likely due to amino acid substitutions at key ACE2-binding sites, e.g. the F486L substitution in RaTG13 weakens hydrophobic interactions with side chain residues L79, M82 and Y83 in ACE2 [15]. Importantly, this implies that generalism and specialism cannot entirely be predicted by phylogenetic relationship and, indeed, a limitation of our study is that we did not look at more viruses to understand this further. Consistent with other studies, the clade II viruses did not use ACE2 as a functional receptor, despite robust Spike protein expression. This can largely be attributed to two deletions in the RBD spanning amino acids 445-449 (region 1) and 473-488 (region 2), relative to SARS-CoV-2, which have been described elsewhere as being indicative of ACE2 tropism [40]. However, substitutions of these regions, or the entire receptor binding motif with SARS-CoV-2, failed to restore ACE2 binding or entry [17, 40], so these are not the only restrictions to ACE2 usage. The clade III viruses RhGB07 and RfGB02 had restricted bat and slightly broader mammalian ACE2 usage, with RhGB07 being able to utilise hACE2, as previously described [10]. Interestingly, although the RBD sequences of RfGB02 and RhGB07 are identical at the RBD/ACE2 binding interface, the former could not use hACE2 - indicating again sequences outside the direct ACE2/RBD interface can influence ACE2 usage [10]. Our structural data for RhGB07 identified that the RhGB07 deletion at position 449 (region 1) potentially disrupts electrostatic interactions with residues 38 and 42 of ACE2, which have also been shown to be important for bat ACE2 receptor tropism [19], which may explain the specialist receptor tropism of this virus. Similar results have been seen for a related clade III virus, PRD-0038, which branches separately from RhGB07 and RfGB02 [14]. Although this virus also uses hACE2 poorly, its host range extends to geographically relevant bat species, *Rh. alcyone*, *Rh. landeri* and specific alleles of *Rh. ferrumequinum* ACE2. One or two amino acid substitutions in Spike – T487W and K482Y – were shown to be sufficient to increase tropism for these bat ACE2s [14]. Conversely, Khosta-2 (another clade III virus) was shown to be able to bind human ACE2 in structural models and infect human cells *in vitro* using full-length Spike, when in the presence of trypsin [41, 42]. Lastly, the clade V Spike Rc-o319 exhibited similarly broad restrictions in bat ACE2 usage and was unable to use human or other mammalian ACE2s. Rc-o319 contains a 10 amino acid deletion in region 2 (481-499), which likely contributes to this specialist receptor usage profile [40]. Of note, viruses from clade IV were not included due to the unavailability of complete sequences at the time of commencing our study. Indeed, one limitation of our approach is that sequences were selected based on data from similar studies [10, 14, 18, 31, 41, 43], however this led to overrepresentation of certain clades. In the future we could use more unbiased criterion to select sequences, as we recently described for the alphacoronaviruses, where Spike libraries were constructed algorithmically to maintain maximum genetic diversity [44]. Nevertheless, other studies have since investigated the tropism of a clade IV virus, RaTG15, which shares 72.6% amino acid identity with SARS-CoV-2 RBD. The RaTG15 RBD bound to *Rh. affinis* and Malayan pangolin ACE2, and full-length Spike pseudoviruses were able to use these receptors for entry, however, no obvious binding or entry with hACE2 was detected [31]. A short deletion present in RaTG15 at amino acid positions 444-447, as well as variation at key ACE2 receptor binding sites (486, 493, 494, 501) is likely to explain this specialism.

Although this study focused on bat sarbecoviruses, related viruses have now been isolated from *Paguma larvata* (palm civet, e.g., SARS CoV SZ3 in China) and *Manius javanica* (pangolin, e.g., BetaCoV/P4L in China), respectively [18]. These hosts have been proposed as potential intermediate reservoirs for SARS-CoV-2, along with the raccoon dog [45], so future host range analyses should include these sarbecoviruses to understand the breadth of species tropism of possible virus intermediates. That being said, the larger unanswered question that remains is to understand what are the environmental selection pressures that maintain receptor generalism for some sarbecoviruses but not others. It is attractive to postulate that the overlapping host-range of multiple bat species in Southeast Asia leads to significant selection pressure for generalism in resident sarbecoviruses (mostly clade 1), ecological pressures that are less prevalent in regions where specialists were isolated. However, the identification of specialists within clade 1b like RaTG13 is indicative of a more complicated and perhaps stochastic pattern.

Another important consideration is to understand the impact of the extensive, ongoing human adaptation of SARS-CoV-2 on its ‘reverse zoonosis’ potential. The oscillating pattern of bat ACE2 usage by SARS-CoV-2 variants was an interesting observation, especially the eventual drift away from bat tropism by later Omicron variants. We identified the Q493R mutation in Omicron BA.1, either by itself or in combination with other RBD-specific mutations (N501Y, E484A and K417N) as being an important determinant of this trend. Q493R has previously been shown to be important in overcoming bat ACE2 restrictions, due to an increased affinity for ACE2 residues 35 and 31, which were described as key determinants for bat ACE2 usage [46], along with ACE2 amino acid positions 27 and 42 [19, 34]. Interestingly, in the BA.5 variant position 493 is reverted (R493Q), and we saw a corresponding loss of *Rh. cornutus* and *Rh. ferrumequinum* ACE2 usage here. Other notable animal ACE2 restrictions observed are for the platyrrhines ACE2s (marmoset and squirrel monkey), which exhibit lower levels of entry compared to catarrhines (lemur and macaque). This is attributable to ACE2 variation at positions 41 and 42 and the loss of important hydrogen bonds (which help binding to positions 446 and 500 in Spike) [47]. Despite shifts in bat ACE2 usage most SARS-CoV-2 variants have maintained their generalist receptor usage with a wider selection of mammal ACE2s. The impact of this generalism is now more widely appreciated than in 2020, as evidenced by white-tailed deer infections in the USA [25, 27], mink in Denmark [37], and hamsters in China [38]. The second objective of our study was to relate this generalism and specialism to antigenicity. For example, the broad host range we observed for many clade I sarbecoviruses raises concerns that these viruses might spillover into human populations. Accordingly, we investigated the breadth of protection provided by COVID-19 derived immunity. For clade I viruses, we observed broad evidence for cross neutralisation by sera from convalescent individuals, regardless of the infecting variant. Indeed immunisation of human ACE2-transgenic hamsters with a trivalent SARS-CoV-2/SHC014/XBB Spike nanoparticle vaccine also exhibited complete protection against challenge with other clade I viruses, WIV-1 and SHC014 [48], supporting our data demonstrating pan-clade I neutralisation from clade I derived convalescent sera. We also noted some evidence of neutralisation against more specialised ACE2 users, e.g. RhGB07 and Rc-o319, with patterns of neutralisation that mirrored those observed for the antigenically distinct SARS-CoV-2 variant BA.1. However, the overall affinity and avidity of Spike for human ACE2 is worth considering and understanding, in relation to these more distantly related viruses. For example, a study examining the receptor tropism of another clade III virus – Khosta 2 – exhibited binding of the RBD of this virus with human ACE2, but viral pseudotypes of expressing the Khosta 2 RBD were not neutralised by SARS-CoV-2 derived mAbs or vaccine sera [42]. Furthermore, we previously showed that RaTG13 Spike-based pseudotypes were neutralised more efficiently than SARS-CoV-2, using sera from COVID-19 vaccinated healthcare workers (with or without evidence of prior infection) [16]. A later cryoEM structure of the RaTG13 Spike revealed a more stable, closed conformation for the trimeric protein with low relative affinity for hACE2, compared to SARS-CoV-2 [49, 50]. Our structural data for the RhGB07 trimer, which was also only found in a three-RBD down closed confirmation, is consistent with a similar pattern. The low overall affinity for human ACE2 of these bat virus RBDs, unlike BA.1 for example, could mean that even low-affinity nAbs are able to displace the RBD from human ACE2. In this context, extending our analysis to include sera from vaccinated individuals in the UK would be of interest to understand whether vaccine-derived immunity can enhance binding and neutralisation of bat sarbecoviruses. Nevertheless, to understand the cross-neutralisation we observed with convalescent sera in more detail, we instead expanded our study to include monoclonal antibodies derived from SARS-CoV-2 breakthrough infections. We identified three high affinity antibodies, which belonged to three different RBD competition groups that were able to bind all sarbecovirus RBDs in our screen (apart from RhGB07), including the non-ACE2 using RacCS203. These mAbs were previously classified by competition ELISAs, using mAbs that bind known epitopes in Spike [39, 51, 52] and further characterised in our study by SPR. In our companion article we determine the molecular mechanisms by which these three mAbs retain broad and potent binding. These data provide promising insights into the regions of Spike that could be targeted to produce pan-sarbecovirus vaccines, or indeed the potential for these monoclonals to be used as pan-sarbecovirus therapeutics.

As discussed above our findings provide promising insights into the role COVID-immunity may play in preventing future sarbecovirus spillover. However, an outstanding question that remains from our research is, ‘are there human ACE2-using configurations for coronavirus Spike RBDs that are entirely antigenically distinct from SARS-CoV-2?’ In simpler terms is there another human-ACE2 using serotype of sarbecoviruses in the wild? One hypothesis, favourable from a public health perspective, is that any ‘specialist’ sarbecovirus that evolves to use human ACE2 (a ‘generalist’) will become antigenically recognisable by the population’s COVID-primed immune response. This would set the bar for spillover higher than anticipated; however, the emergence of very long branch SARS-CoV-2 variants, like BA.1 and BA.2.86, clearly highlights a great deal of plasticity in the RBD in binding human ACE2 very efficiently, while still navigating significant immune selection pressure. In addition, viruses like RhGB07 – specialists that can still partially use hACE2 – provide clear evidence that hACE2 usage can arises relatively early on the spectrum of specialism to generalism.

## Supplemental Tables and Figures

**Supplementary Figure 1:**
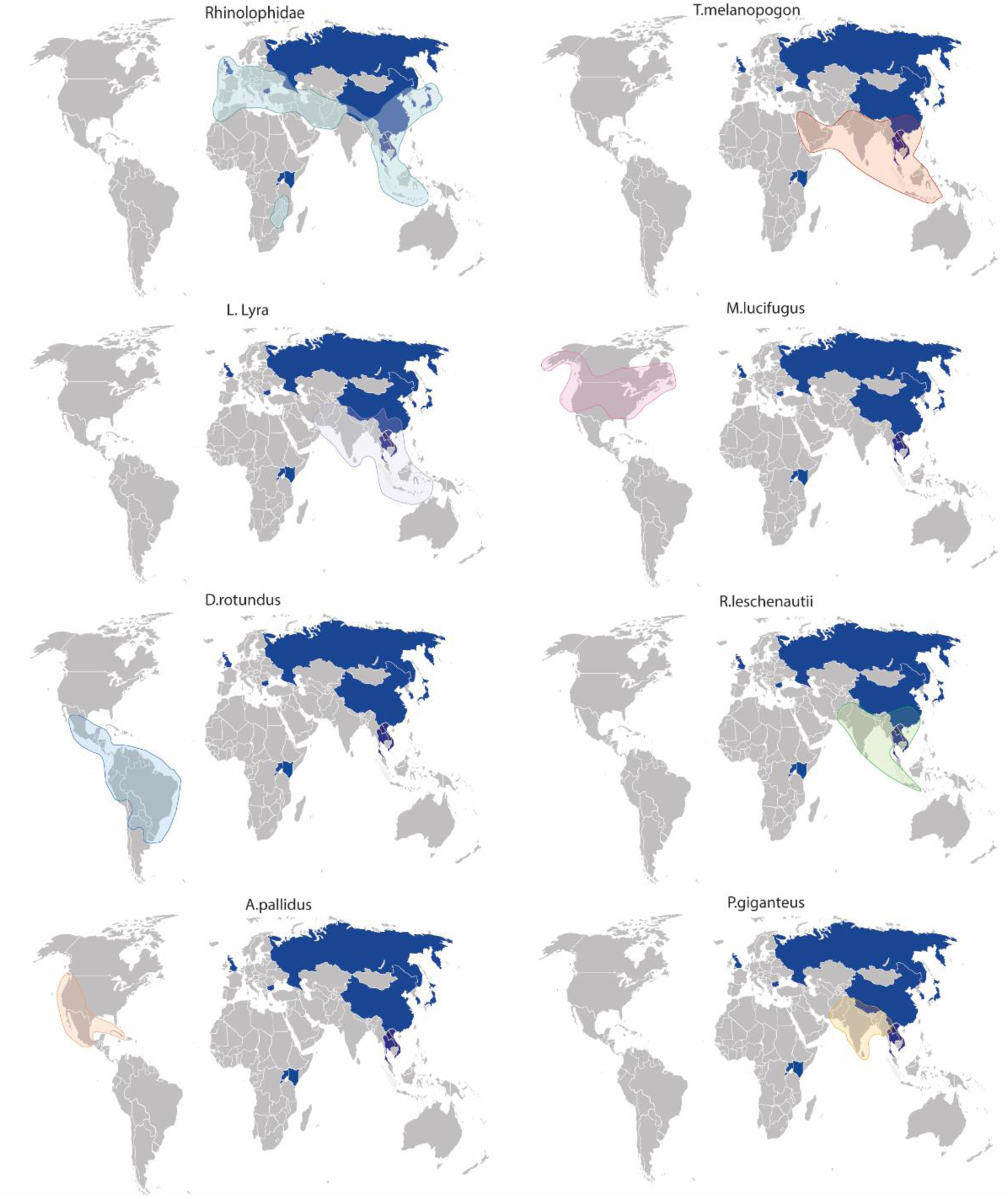
Geographic range of bat species investigated in this study. The geographic location in which the bat sarbecoviruses analysed in this study have been isolated is coloured in blue on the world map, with separate maps depicting the range the bat species investigated in this study span. *Rhinolophidae* species have been grouped together and span across Europe, Asia and some parts of southern Africa. The *T.melanopogon, P. giganeteus, L. Lyra* and *R. leschenautii* species all span across southeast Asia, in particular overlapping in regions of India and parts of China. *The M.lucifugus, D. rotundus* and *A.pallidus* bat species can be found nesting in the Americas.

**Supplementary Figure 2:**
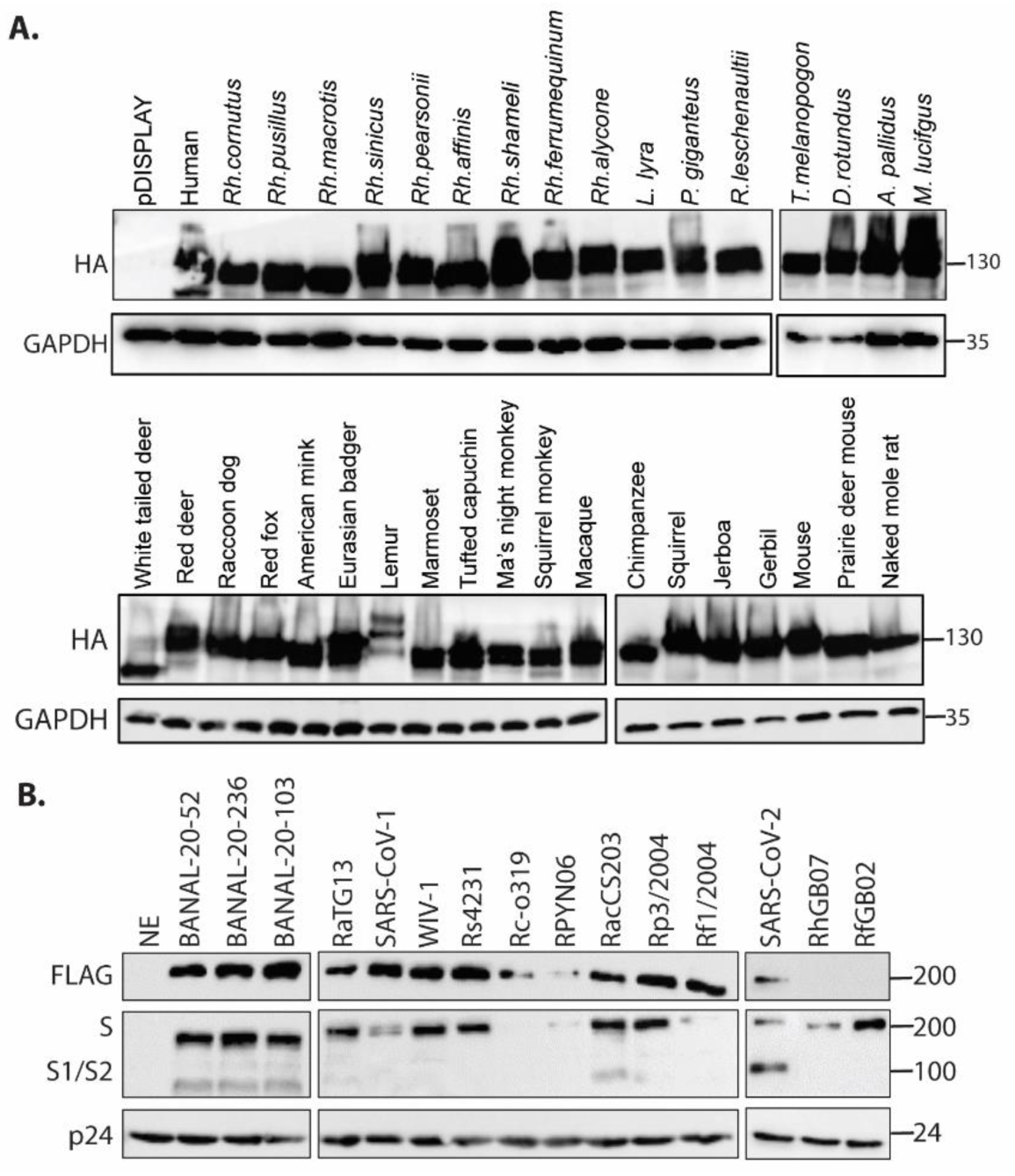
Protein expression of ACE2s and sarbecovirus Spikes. Expression of tagged ACE2 proteins was carried out in lysates from BHK-21 cells transfected with ACE2 constructs and probed for HA expression by Western blotting. Sarbecovirus Spikes were used to generate lentivral-based pseudotypes, and the viral supernatant purified by ultra-centrifugation under a 20% sucrose cushion. Purified viruses were probed with an anti-FLAG antibody (except RhGB07 and RfGB02, which are untagged), an anti-Spike antibody (S, full-length Spike; S1/S2, cleaved variant) and the lentivirus internal control, anti-p24 antibody.

**Supplementary Figure 3:**
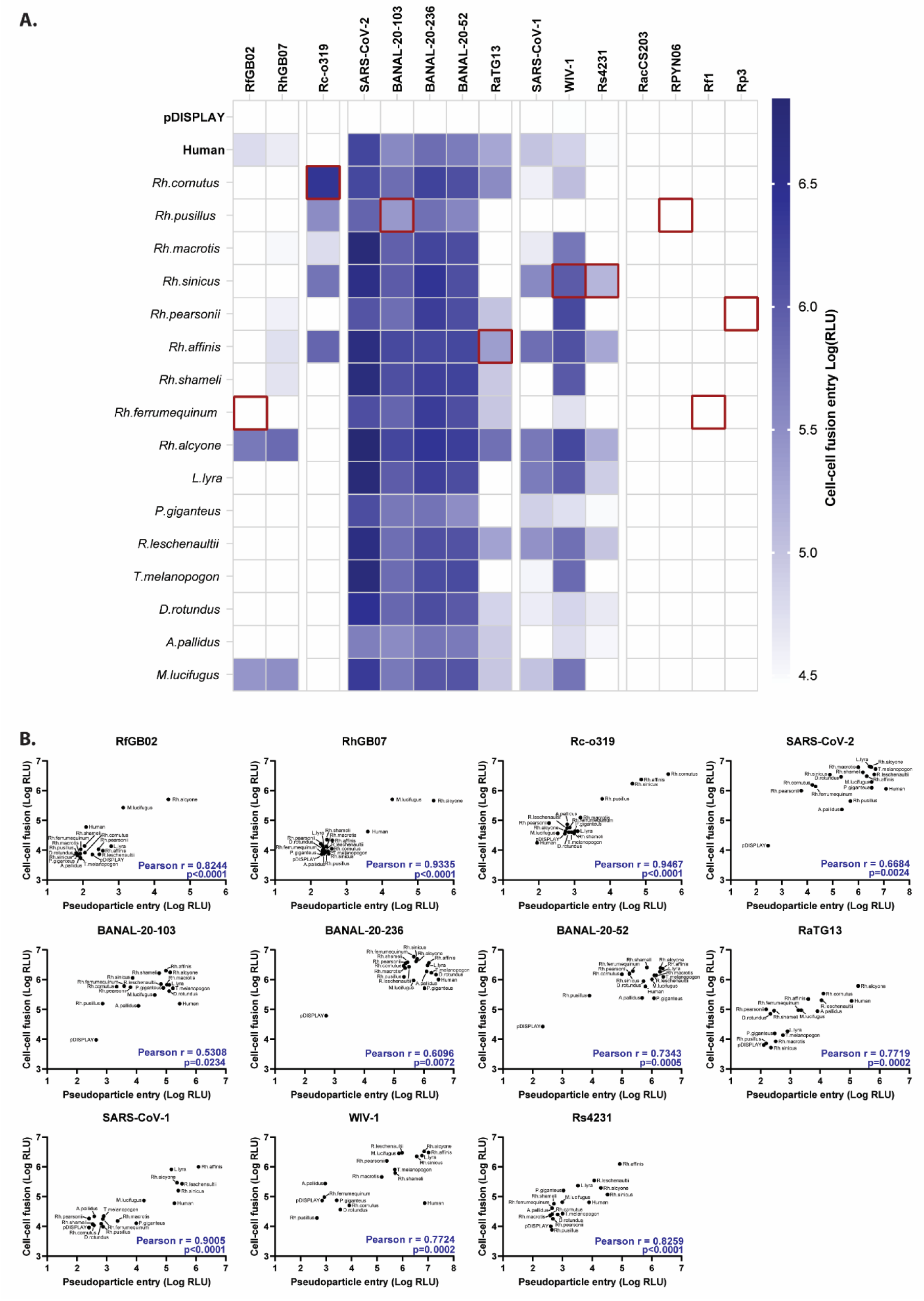
Entry of bat sarbecoviruses by cell-cell fusion. **(A)** Heatmap showing virus entry by cell-cell fusion. HEK293T cells stably expressing half of a split GFP-Renilla (RLuc-GFP 1-7) luciferase reporter were transfected with bat sarbecovirus Spikes, and co-cultured with BHK-21 cells expressing the other half of the split reporter (RLuc-GFP 8-11), transfected with bat ACE2s. Productive cell-cell fusion results in reconstitution of the split reporter, from which luciferase signals can be measured. Data is shown as log (RLU) and is the average of three separate experiments. A pDISPLAY vector control was included to set the baseline value to account for background signals. The red boxes indicate the predicted cognate bat receptor for each bat sarbecovirus, where known. **(B)** XY scatter plots for each of the viruses assessed for their receptor usage profile, correlating the log (RLU) of pseudoparticle entry (X) against cell-cell fusion entry (Y), excluding clade II viruses that do not use ACE2 as their entry receptor (RfGB02: r= 0.8244, p<0.0001; RhGB07: r=0.9335, p<0.0001; Rc-o319: r=0.9467, p<0.0001; SARS-CoV-2: r = 0.6684, p=0.0024; BANAL-20-103: r=0.5308, p=0.0234; BANAL-20-236: r=0.6096, p=0.0072; BANAL-20-52: r=0.7343, p=0.0005; RaTG13: r=0.7719, p=0.0002; SARS-CoV-1: r=0.9005, p<0.0001; WIV-1: r=0.7724, p=0.0002; Rs4231: r=0.8259, p<0.0001). Each data point is labelled with the ACE2 corresponding ACE2 receptor and a Pearson’s correlation calculated with 95% confidence intervals. The Pearson r value and p value are both documented.

**Supplementary Figure 4:**
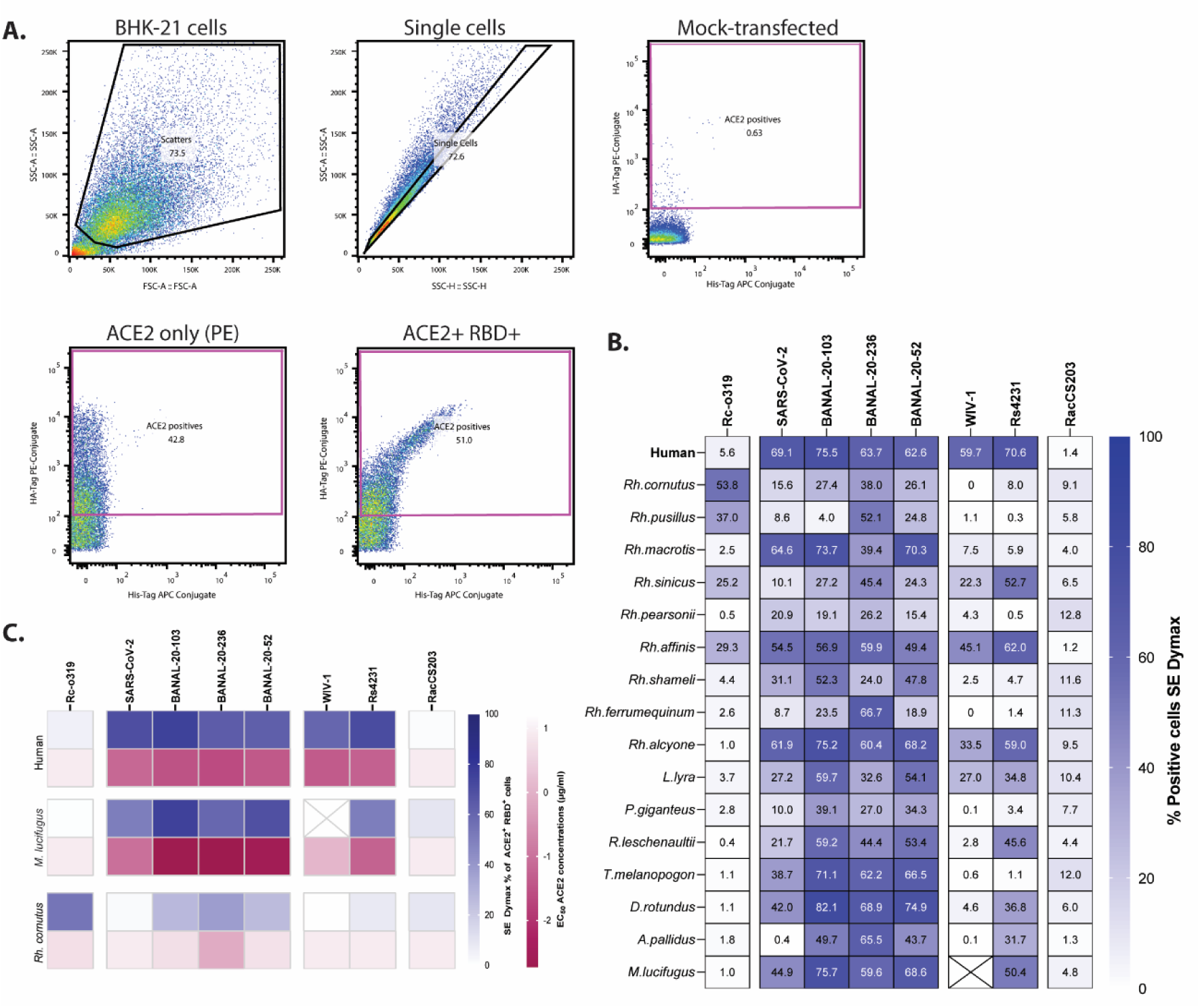
Gating strategy and analysis for flow cytometry of ACE2-expressing constructs bound to sarbecovirus RBDs. **(A)** BHK-21 cells were transfected with HA-tagged bat ACE2-expressing constructs **(S2 Table)** and baited with His tagged-purified RBD proteins for bat sarbecoviruses. Cells were stained with anti-HA PE conjugated antibody (ACE2) and anti-His APC conjugated antibody (RBD). Live and singlet BHK-21 cells were gated as PE+ APC+ (ACE2+RBD+), relative to mock-transfected cells (pDISPLAY). **(B)** The SE% Dymax metric was used to calculate the percentage of ACE2+RBD+ cells, shown for all RBDs tested with our library of bat ACE2 and human ACE2. Representative datasets are shown for mock-transfected cells and raccoon dog ACE2 with SARS-CoV-2 RBD, with their respective SE Dymax % positive value noted. **(C)** Heatmap showing the binding of purified bat sarbecovirus proteins to ACE2s of human, *M. lucifugus Rh. cornutus* by ELISA (pink, EC_50_ µg/ml) or flow cytometry (blue, % ACE2^+^RBD^+^ cells). Data is representative of a mean of 3 separate experiments. A cross in a box indicates this data was not obtained.

**Supplementary Figure 5:**
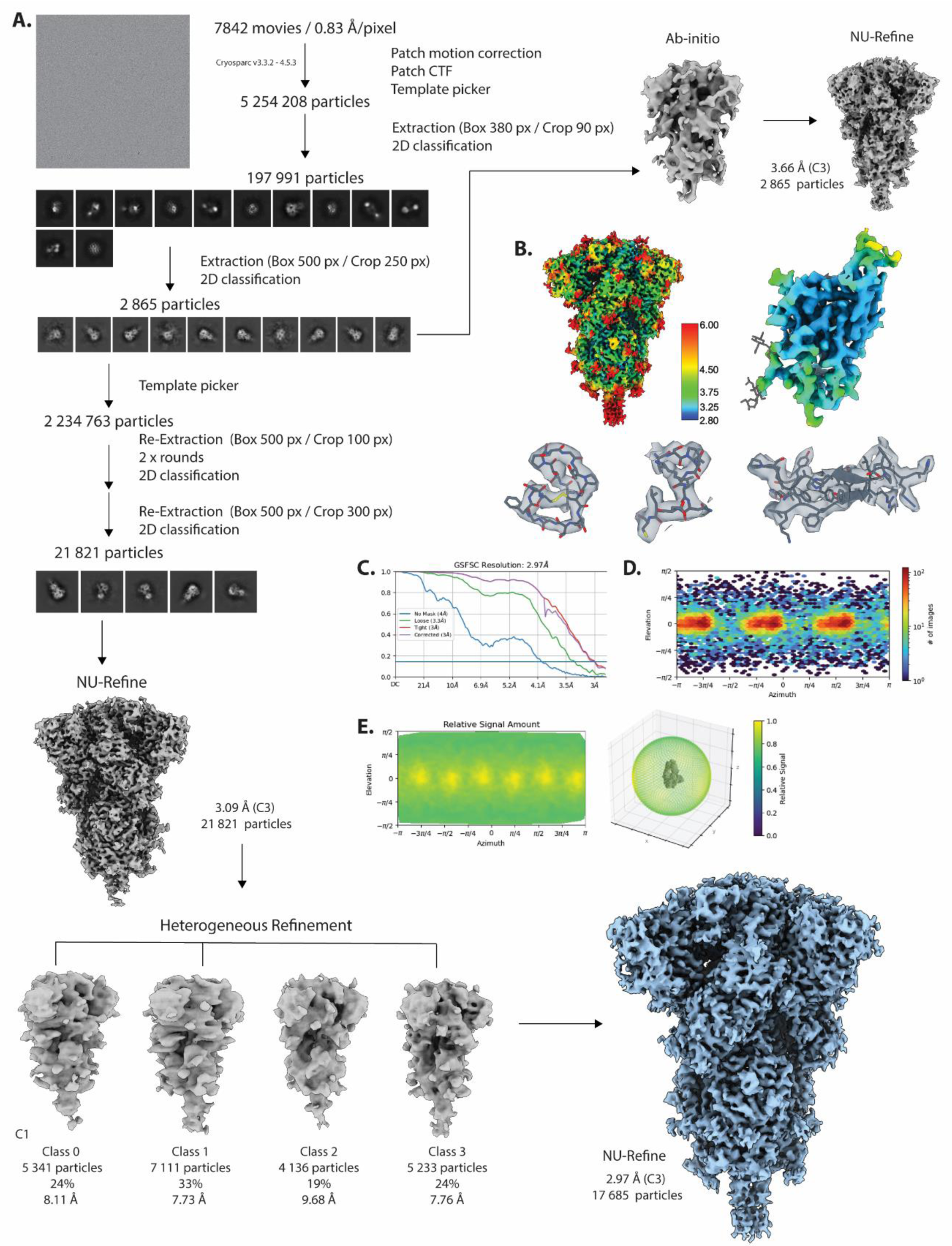
Cryo-EM Processing Workflow. **(A)** Processing workflow of for the determination of the RhGB07 full-length trimeric spike. **(B)** Cryo-EM density coloured by local-resolution of the full-length trimer and of the isolated RBD. Selected regions represented with their cryo-EM density is also shown. **(C)** FSC curve showing a determined resolution for the full-length spike of 2.97 Å **(D)** Viewing direction distribution of particles used in the reconstruction. **(E)** Relative Signal vs Viewing direction plot of particles. Cryo-EM data collection, refinement and validation statistics can be sound in **S3 Table**.

**Supplementary Figure 6:**
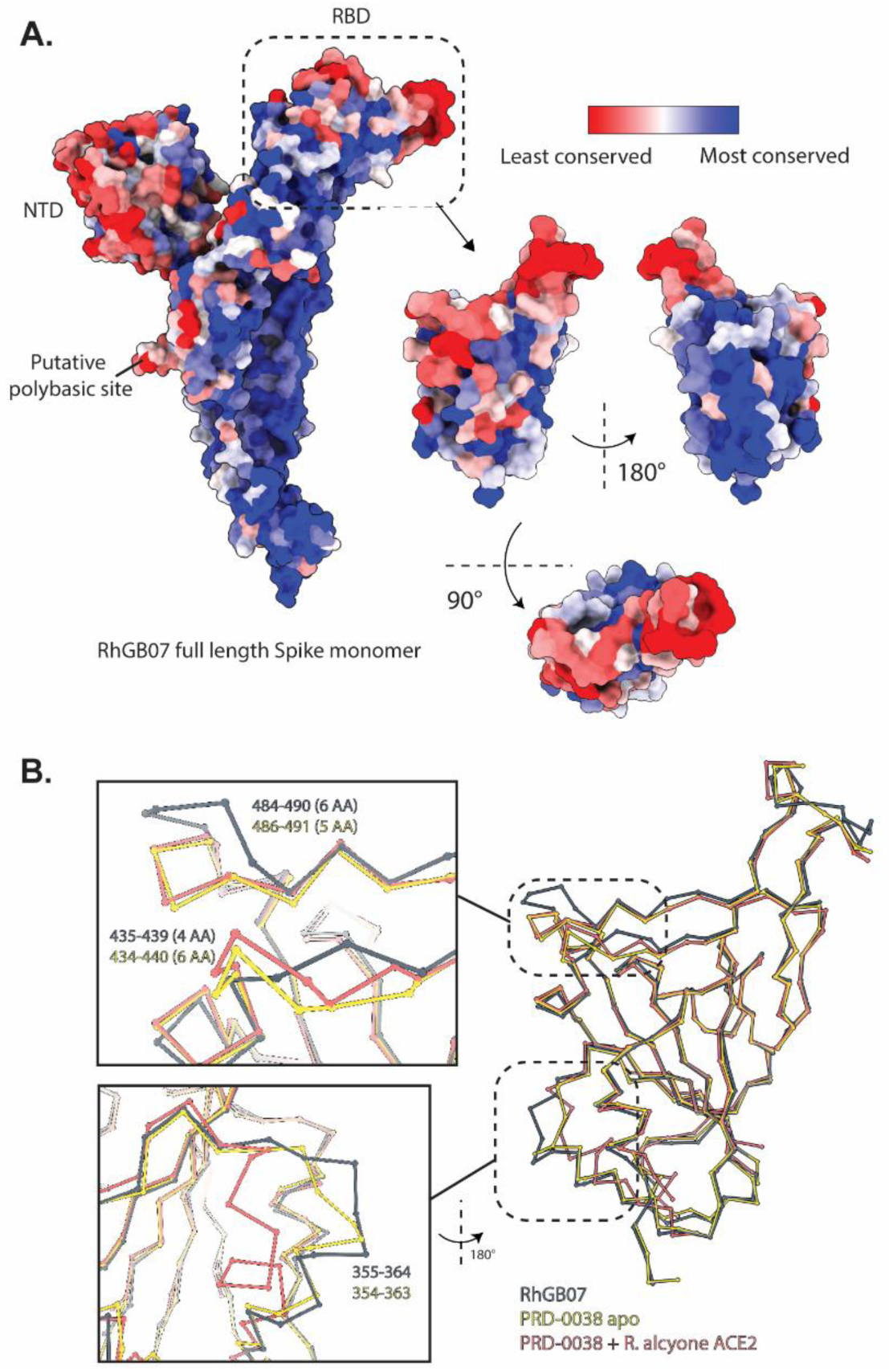
Sequence conservation of the full-length RhGB07 spike. **(A)** Sequence conservation of selected sarbecoviruses were mapped on to the RhGB07 spike. Areas least conserved are in red and most conserved are blue. The least conserved regions are at the top of the RBD, the NTD and close to a putative polybasic site. **(B)** The C-alpha trace of the RBD of RhGB07 rendered as balls-and-sticks superimposed with the RBD of PRD-0038 apo or PRD-0039 bound to R. Alcyone ACE2. The insets show a zoomed-in view of two regions of the RBD that show the greatest changes in the position of the C-alpha of the RBD. The region containing the ins1-loop and delta4-loop coincides with putative ACE2-interacting residues based on mapping the equivalent residues.

**Supplementary Figure 7:**
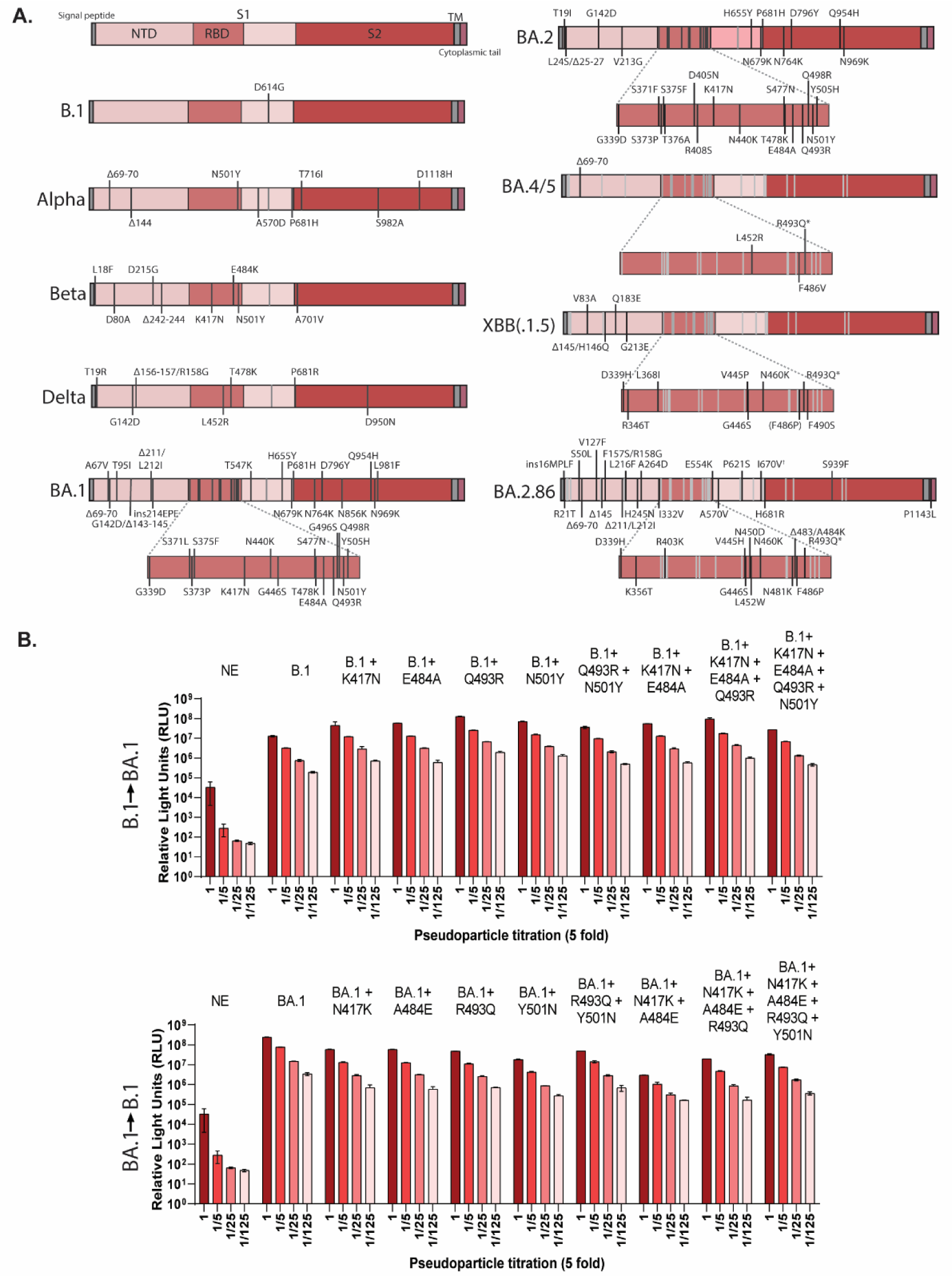
SARS-CoV-2 variants entry and binding with a library of bat ACE2. **(A)** Schematic of the Spike proteins for B.1, Alpha, Delta, Omicron BA.1, Omicron BA.2 and its subvariants BA.4/5, XBB, XBB1.5 and BA.2.86. Key regions of the N-terminal domain (NTD), receptor binding domain (RBD), S1 and S2 are highlighted and the mutations that arose in these variants have been annotated. The BA.4 and BA.5 Spike is identical, and the difference in XBB1.5 compared to XBB is denoted by an asterisk, which is a reversion of the mutation at position 494 (R493Q). **(B)** Infection of HEK293T cells stably expressing hACE2 with SARS-CoV-2, WT and BA.1 pseudoparticles, along with individual mutants generated of these variants at positions 417, 484, 493 and 501, either singly or in combination. Virus was titrated 5-fold on cells and Firefly luciferase signals measured; a negative control (non-enveloped, NE) was also included.

**Supplementary Figure 8:**
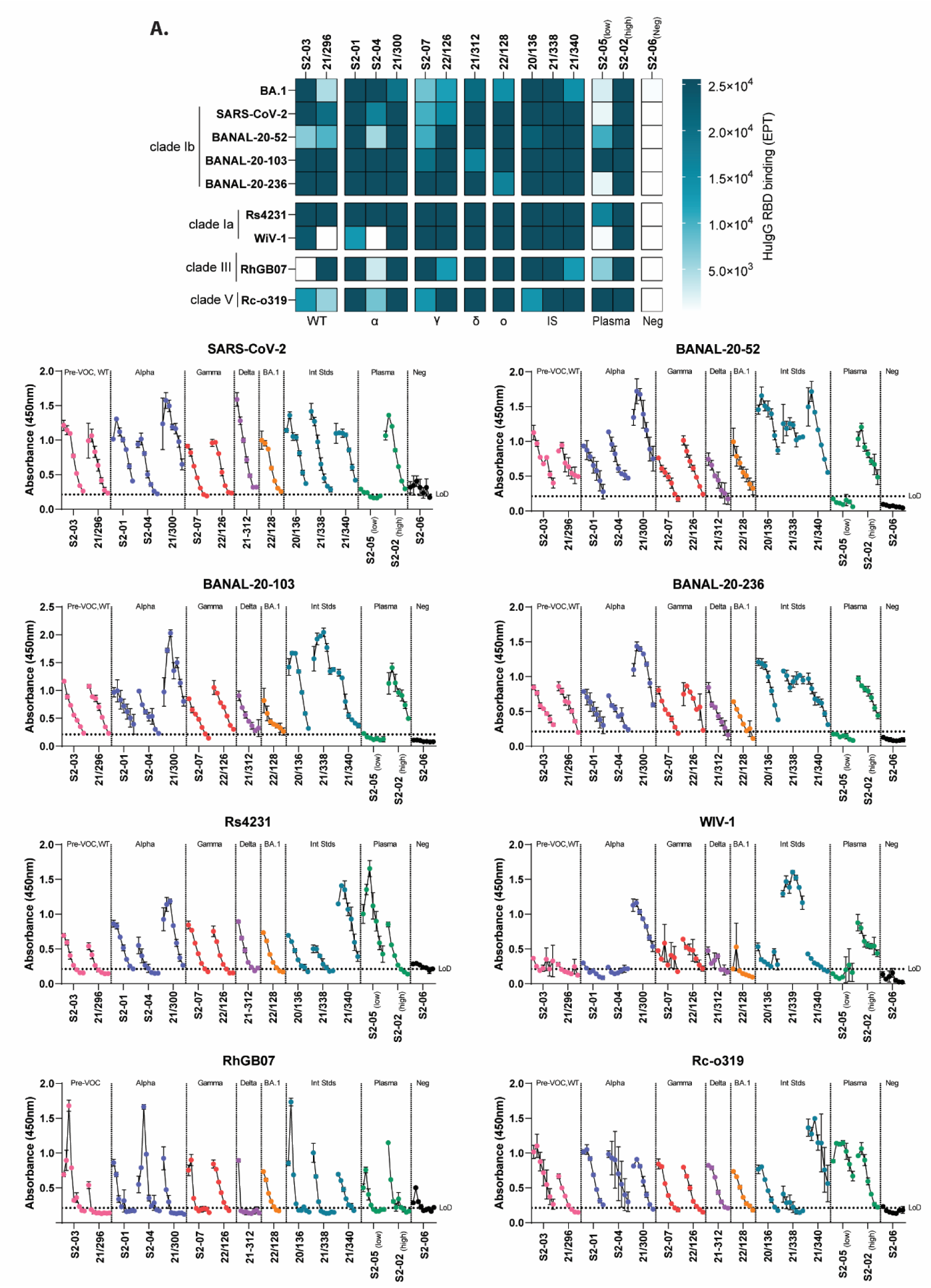
Binding of convalescent sera from SARS-CoV-2 infections with sarbecovirus RBDs. **(A)** Heatmap of calculated human IgG binding end-point titres and **(B)** absorbance binding curves for convalescent sera (details in Figure 5A) with SARS-CoV-2, BANAL-20-52, BANAL-20-103, BANAL-20-236, Rs4231, WIV-1, RhGB07 and Rc-o319 RBDs. Boxes with crosses indicate that sample was not tested. Dotted lines represent the limit of detection for the assay (mean + 3SD of no sera control well), with error bars shown for replicate values (mean + SD).

**Supplementary Figure 9:**
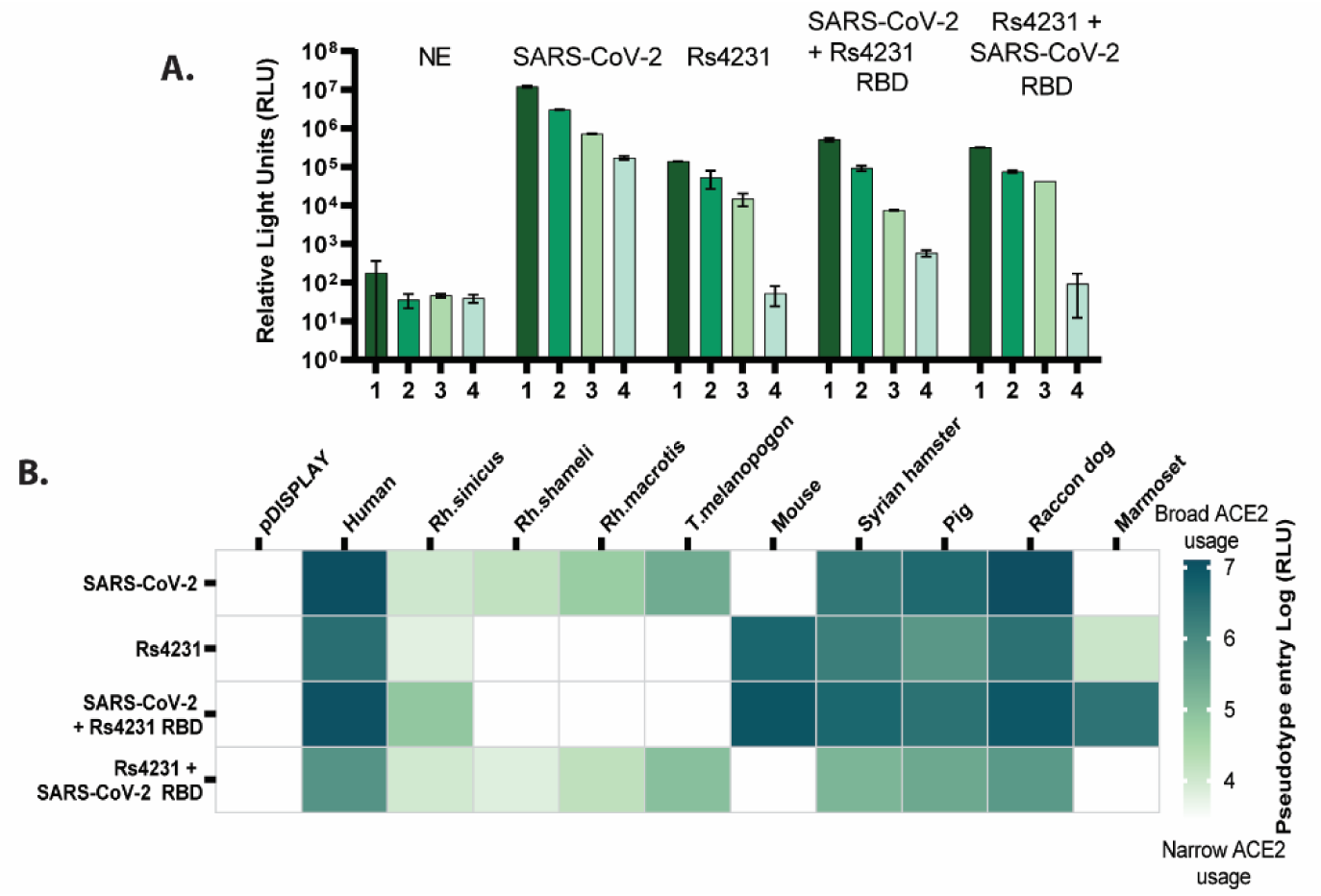
RBD Chimeras of SARS-CoV-2 and Rs4231 show a switch in ACE2 entry phenotype. **(A)** Infection of HEK293T cells stably expressing hACE2 with SARS-CoV-2, Rs4231 or chimeric pseudoparticles (Rs4231 + SARS-CoV-2 RBD, SARS-CoV-2 + Rs4231 RBD) Virus was titrated 5-fold on cells and Firefly luciferase signals measured; a negative control (non-enveloped, NE) was also included. **(B)** Pseudoparticle entry assay using SARS-CoV-2 and Rs4231 chimeras, on a subset of ACE2s that had striking entry profile in the initial screen (human, *Rh. sinicus*, *Rh. shameli, Rh. macrotis, T. melanopogan,* mouse, Syrian hamster, pig, raccoon dog and marmoset ACE2). The heatmap shows log(RLU) of psuedotype entry, with pDISPLAY included as a control to establish background luciferase signal.

**Supplementary Figure 10:**
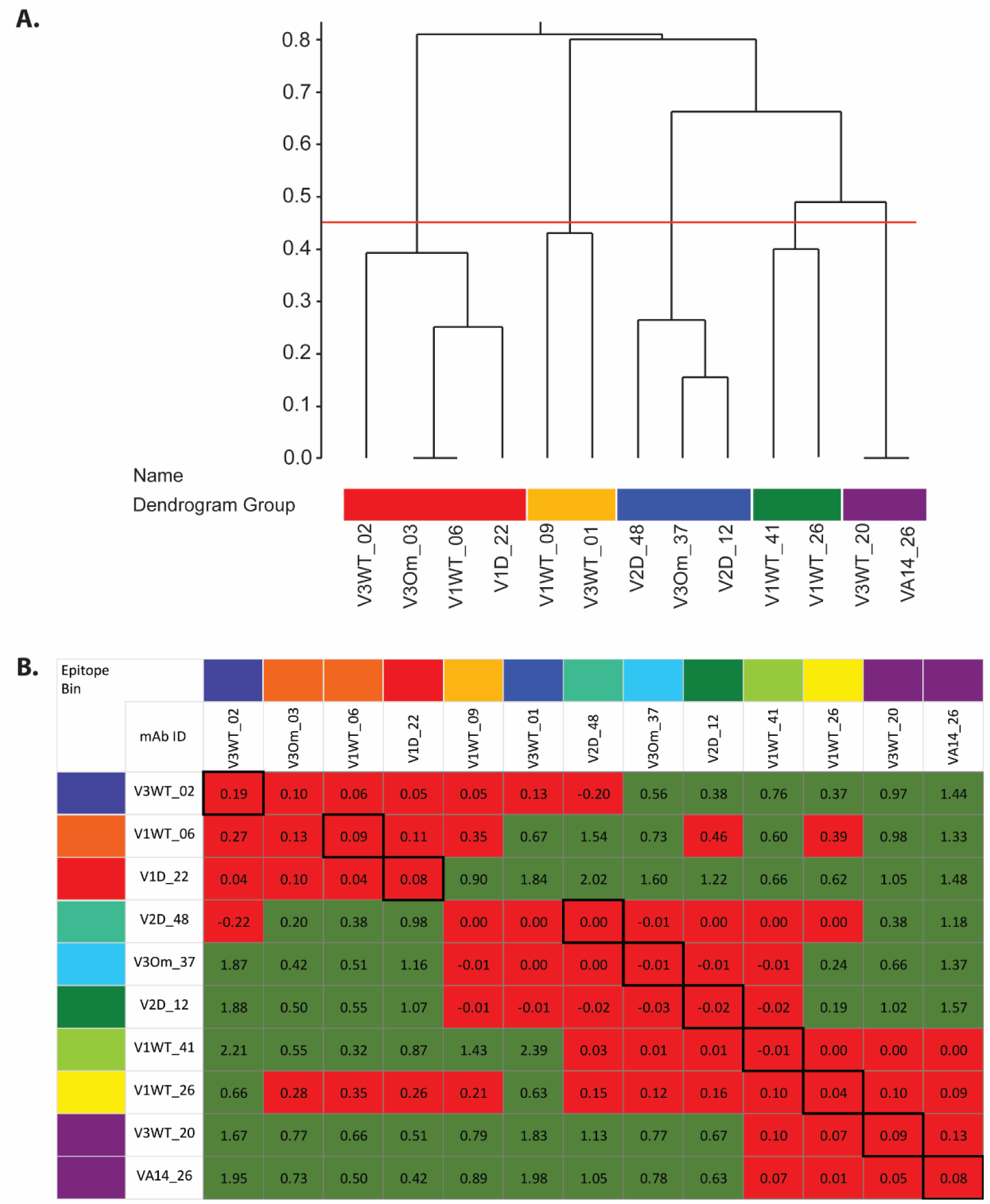
Binding dendrogram of SARS-CoV-2 monoclonal antibodies analysed by epitope binning. **(A)** Dendrogram represents the relatedness of the epitopes of the antibodies. The cut-off for creating antibody communities is shown by the red line at 0.45. **(B)** Heatmap of antibody blocking and sandwiching interactions. Red boxes indicate blocking interactions, green indicates sandwiching interactions. Self-self blocking is indicated by red boxes with thicker black outlines. Antibodies immobilised as ligands are listed on the left and analytes injected are listed across the top. Where a monoclonal antibody failed to bind as a ligand, it has been excluded.

**Supplementary Figure 11:**
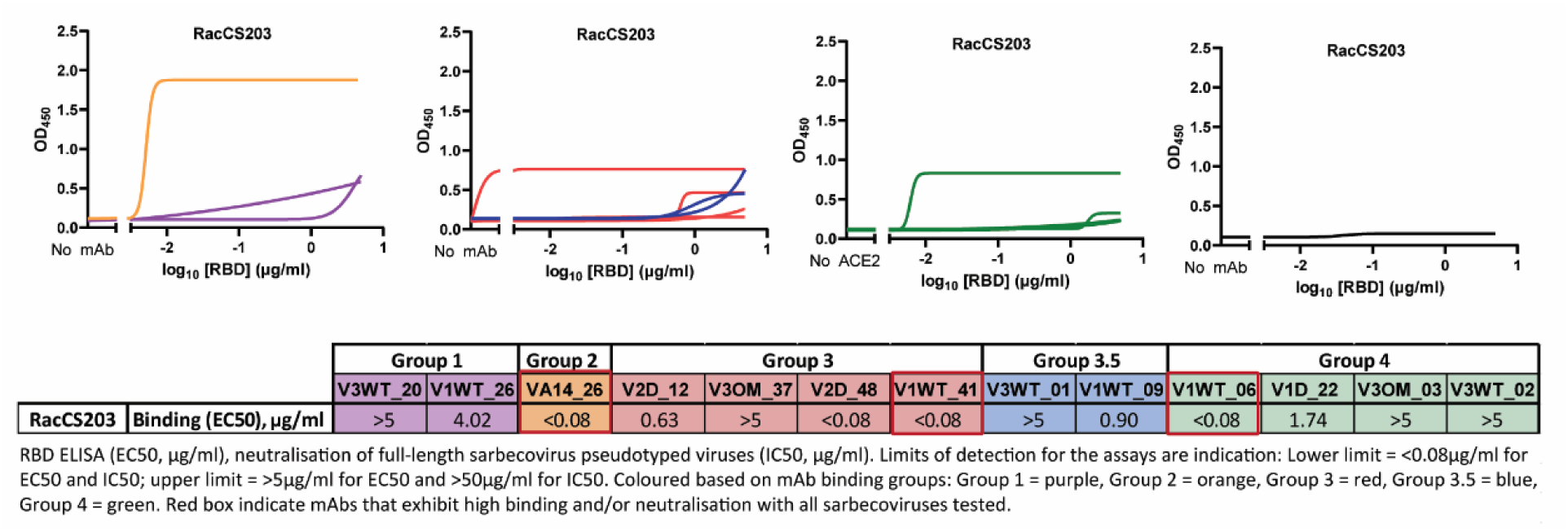
Binding of clade II virsus, RacCS2013 RBD with monoclonal antibodies isolated from SARS-CoV-2 breakthrough infections. **(A)** Each plot shows a single mAb binding, coloured by its designated antibody competition group (Group 1 = purple, Group 2 = orange, Group 3 = red, Group 3.5 = blue, Group 4 = green, Group 8 = non-RBD, NTD-specific) in an ELISA using purified RacCS203 RBD. The curves were calculated using a least squares regression non-linear fit. Binding (EC_50_, µg/ml) titres are shown for each mAb against RacCS203, with limits of detection indicated (lower limit: <0.08g/ml upper limit: 5µg/ml). Red boxes indicate mAbs that exhibit high binding with all sarbecoviruses tested.

**S1 Table:**
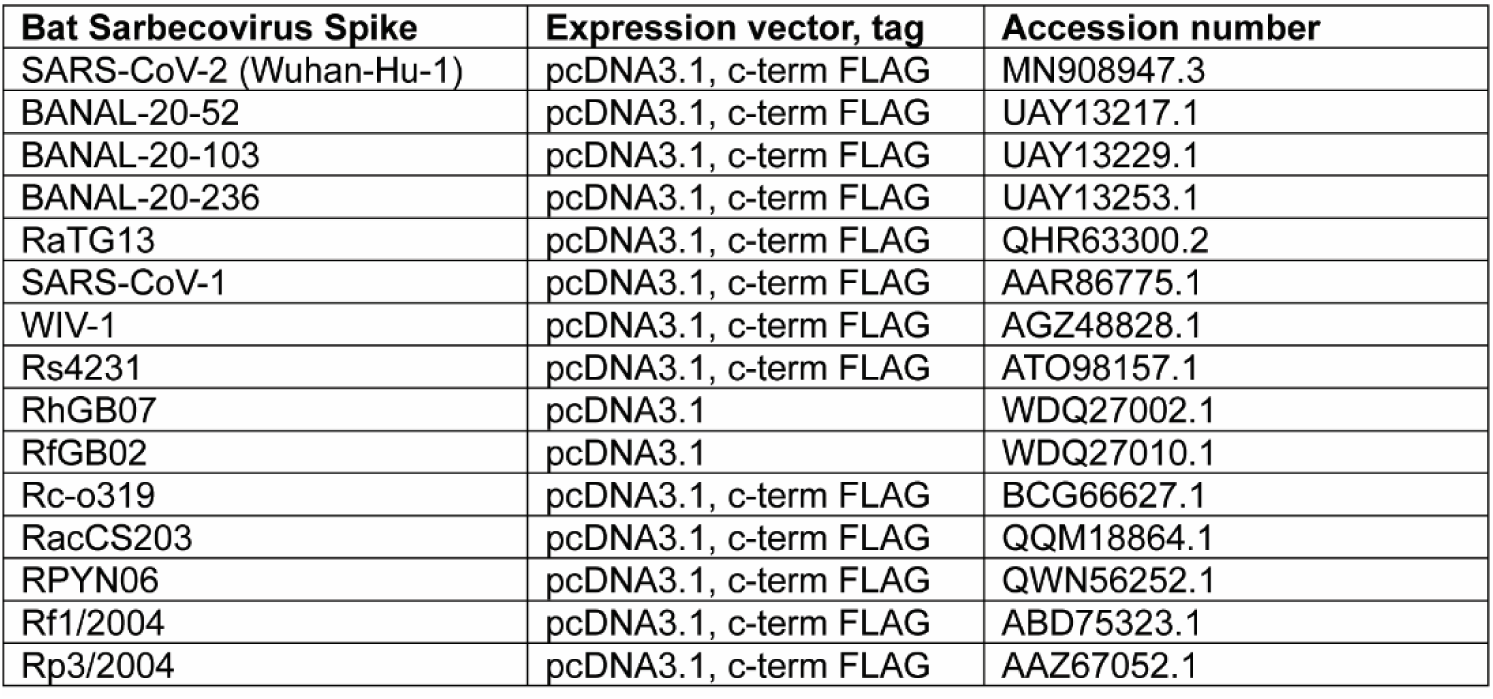
Bat Sarbecovirus Spike constructs used in this study for receptor screens and neutralisation assays.

**S2 Table:**
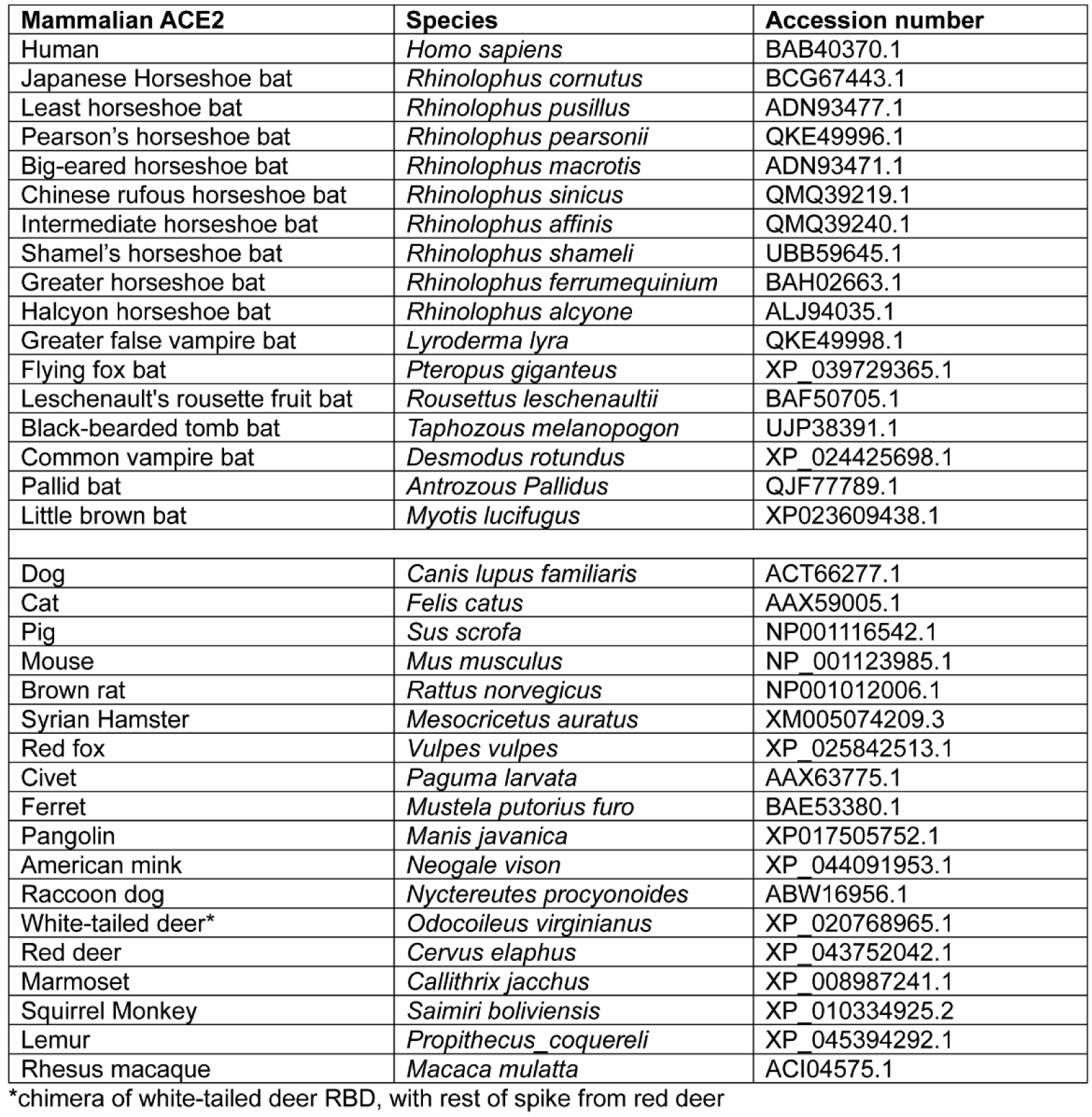
codon optimised ACE-2 expression plasmids used in this study for receptor screens.

| Mammalian ACE2 | Species | Accession number |
| --- | --- | --- |
| Human | *Homo sapiens* | BAB40370.1 |
| Japanese Horseshoe bat | *Rhinolophus cornutus* | BCG67443.1 |
| Least horseshoe bat | *Rhinolophus pusillus* | ADN93477.1 |
| Pearson's horseshoe bat | *Rhinolophus pearsonii* | QKE49996.1 |
| Big-eared horseshoe bat | *Rhinolophus macrotis* | ADN93471.1 |
| Chinese rufous horseshoe bat | *Rhinolophus sinicus* | QMQ39219.1 |
| Intermediate horseshoe bat | *Rhinolophus affinis* | QMQ39240.1 |
| Shamel's horseshoe bat | *Rhinolophus shameli* | UBB59645.1 |
| Greater horseshoe bat | *Rhinolophus ferrumequinum* | BAH02663.1 |
| Halcyon horseshoe bat | *Rhinolophus alcyone* | ALJ94035.1 |
| Greater false vampire bat | *Lyroderma lyra* | QKE49998.1 |
| Flying fox bat | *Pteropus giganteus* | XP_039729365.1 |
| Leschenault's rousette fruit bat | *Rousettus leschenaultii* | BAF50705.1 |
| Black-bearded tomb bat | *Taphozous melanopogon* | UJP38391.1 |
| Common vampire bat | *Desmodus rotundus* | XP_024425698.1 |
| Pallid bat | *Antrozous Pallidus* | QJF77789.1 |
| Little brown bat | *Myotis lucifugus* | XP023609438.1 |
| Dog | *Canis lupus familiaris* | ACT66277.1 |
| Cat | *Felis catus* | AAX59005.1 |
| Pig | *Sus scrofa* | NP001116542.1 |
| Mouse | *Mus musculus* | NP_001123985.1 |
| Brown rat | *Rattus norvegicus* | NP001012006.1 |
| Syrian Hamster | *Mesocricetus auratus* | XM005074209.3 |
| Red fox | *Vulpes vulpes* | XP_025842513.1 |
| Civet | *Paguma larvata* | AAX63775.1 |
| Ferret | *Mustela putorius furo* | BAE53380.1 |
| Pangolin | *Manis javanica* | XP017505752.1 |
| American mink | *Neogale vison* | XP_044091953.1 |
| Raccoon dog | *Nyctereutes procyonoides* | ABW16956.1 |
| White-tailed deer* | *Odocoileus virginianus* | XP_020768965.1 |
| Red deer | *Cervus elaphus* | XP_043752042.1 |
| Marmoset | *Callithrix jacchus* | XP_008987241.1 |
| Squirrel Monkey | *Saimiri boliviensis* | XP_010334925.2 |
| Lemur | *Propithecus coquereli* | XP_045394292.1 |
| Rhesus macaque | *Macaca mulatta* | ACI04575.1 |

\*chimera of white-tailed deer RBD, with rest of spike from red deer

**S3 Table:** Cryo-EM data collection, refinement, and validation statistics.

| Data collection and processing | Structure of the RhGB07 Prefusion, Closed-state trimeric spike protein (EMDB: 53938) (PDB: **pdb_00009RDT**) |
| --- | --- |
| Magnification | 96kx |
| Voltage (kV) | 300 |
| Microscope | TFS Titan G4 |
| Electron exposure (e-/Å<sup>2</sup>) | 50 |
| Defocus range (μm) | -1.0 to -2.5 |
| Pixel size (Å) | 0.83 |
| Symmetry imposed | C3 |
| Initial particle images (no.) | 5 254 208 |
| Final particle images (no.) | 17 685 |
| Map resolution (Å) | 2.97 |
| FSC threshold | 0.143 |
| **Refinement** |  |
| Map sharpening B factor (Å<sup>2</sup>) | -51.2 |
| Model resolution (Å) | 3.4 |
| FSC threshold | 0.5 |
| Model composition |  |
| Non-hydrogen atoms | 27102 |
| Protein residues | 3273 |
| Water | 0 |
| Ligands | NAG:99 BMA:21 MAN: 6 |
| B factors (Å<sup>2</sup>) (min/max/mean) |  |
| Protein | 49.24/199.09/106.88 |
| Ligands | 36.50/217.27/115.22 |
| R.m.s. deviations (# > 4σ) |  |
| Bond lengths (Å) | 0.002 (0) |
| Bond angles (°) | 0.560 (3) |
| Validation |  |
| MolProbity score | 1.67 |
| Clash score | 4.73 |
| Poor rotamers (%) | 1.48 |
| Ramachandran plot |  |
| Favored (%) | 95.76 |
| Allowed (%) | 4.24 |
| Disallowed (%) | 0.00 |

**S4 Table:** Binding of bat sarbecovirus RBDs and neutralisation of bat sarbecovirus Spike pseudotypes with monoclonal antibodies.

|  |  | Group 1 | Group 2 | Group 3 |  | Group 3.5 |  | Group 4 |  |
| --- | --- | --- | --- | --- | --- | --- | --- | --- | --- |
|  |  | V3WT_20 | VA14_26 | V2D_12 | V1WT_41 | V3WT_01 | V1WT_09 | V1WT_06 | V3OM_03 |
| SARS-CoV-2 | Binding (EC<sub>50</sub>, µg/ml) | <0.08 | <0.08 | <0.08 | <0.08 | <0.08 | <0.08 | <0.08 | 0.14 |
|  | Affinity (KD) | 4.35x10<sup>-10</sup> | 2.90x10<sup>-10</sup> | 4.38x10<sup>-10</sup> | 1.29x10<sup>-09</sup> | 2.2x10<sup>-09</sup> | 3.42x10<sup>-09</sup> | 1.02x10<sup>-10</sup> | 1.20x10<sup>-10</sup> |
|  | Neutralisation (IC<sub>50</sub>, µg/ml) | 0.28 | 0.12 | <0.08 | 0.15 | <0.08 | 0.10 | 0.76 | 0.42 |
| BANAL-20-52 | Binding (EC<sub>50</sub>, µg/ml) | <0.08 | <0.08 | <0.08 | <0.08 | <0.08 | 0.08 | <0.08 | 0.15 |
|  | Affinity (KD) | 5.20x10<sup>-08</sup> | 9.25x10<sup>-10</sup> | 3.60x10<sup>-10</sup> | 6.05x10<sup>-10</sup> | 2.20x10<sup>-09</sup> | 1.63x10<sup>-08</sup> | 8.52x10<sup>-11</sup> | 3.40x10<sup>-09</sup> |
|  | Neutralisation (IC<sub>50</sub>, µg/ml) | >50 | <0.08 | <0.08 | <0.08 | <0.08 | 0.18 | 1.22 | 0.50 |
| Rs4231 | Binding (EC<sub>50</sub>, µg/ml) | 0.71 | <0.08 | >5 | <0.08 | >5 | >5 | <0.08 | 5.00 |
|  | Affinity (KD) | 8.48x10<sup>-09</sup> | 5.73x10<sup>-10</sup> | - | 8.98x10<sup>-10</sup> | 2.25x10<sup>-08</sup> | - | 5.05x10<sup>-10</sup> | 2.90x10<sup>-05</sup> |
|  | Neutralisation (IC<sub>50</sub>, µg/ml) | 37.64 | <0.08 | >50 | <0.08 | >50 | >50 | <0.08 | >50 |
| Rc-o319 | Binding (EC<sub>50</sub>, µg/ml) | 1.84 | 0.32 | >5 | <0.08 | 2.40 | 1.71 | <0.08 | >5 |
|  | Affinity (KD) | - | 1.38x10<sup>-07</sup> | - | 1.40x10<sup>-08</sup> | 1.62x10<sup>-06</sup> | - | 4.13x10<sup>-09</sup> | 1.48x10<sup>-06</sup> |
|  | Neutralisation (IC<sub>50</sub>, µg/ml) | >50 | >50 | >50 | 15.90 | >50 | >50 | 11.60 | >50 |
| RhGB07 | Binding (EC<sub>50</sub>, µg/ml) | <0.08 | >5 | <0.08 | <0.08 | 0.34 | 1.69 | <0.08 | >5 |
|  | Affinity (KD) | - | - | - | 4.48x10<sup>-08</sup> | 9.15x10<sup>-09</sup> | 1.17x10<sup>-08</sup> | 4.85x10<sup>-11</sup> | - |

Sarbecovirus RBD ELISA (EC<sub>50</sub>, µg/ml) SPR binding kinetics dissociation constant using sarbecovirus RBD (KD, M), and neutralisation of full-length sarbecovirus Spike pseudotyped viruses (IC<sub>50</sub>, µg/ml). Limits of detection for the assays are indicated (Lower limit = <0.08µg/ml for EC<sub>50</sub> and IC<sub>50</sub>; upper limit = >5µg/ml for EC<sub>50</sub> and >50µg/ml for IC<sub>50</sub>). No value for KD (-) indicates no binding of RBD and mAb by SPR. Boxes are coloured based on mAb binding competition groups: Group 1 = purple, Group 2 orange, Group 3 = red, Group 3.5 = blue, Group 4 = green. Red box indicates mAbs the exhibit high binding and/or neutralisation with all sarbecoviruses tested.

